# Intranasal sarbecovirus vaccine booster elicits cross-clade, durable and protective systemic and mucosal immunity

**DOI:** 10.1101/2025.04.04.647326

**Authors:** Nicholas You Zhi Cheang, Wee Chee Yap, Kirsteen McInnes Tullett, Xinlei Qian, Peck Szee Tan, Kiren Purushotorman, Wan Yi Tan, Shirley Yun Yan Mah, Paul Anthony Macary, Chee Wah Tan, Mireille Hanna Lahoud, Sylvie Alonso

## Abstract

Short-lived, clade-specific immune responses with limited mucosal priming are limitations faced by current COVID-19 mRNA vaccines against sarbecoviruses. We have developed a nasal booster vaccine candidate that induced robust and sustained, cross-clade, systemic and mucosal protective immunity. Two recombinant Clec9A-specific monoclonal antibodies fused to the Receptor Binding Domain (RBD) from Omicron XBB.1.5 and SARS-CoV-1, respectively were generated. In Comirnaty mRNA-vaccinated mice, boosting with each individual Clec9A-RBD construct induced immune responses that either were limited in breadth or waned over time; while boosting with both constructs combined (Clec9A^OMNI^) elicited robust cross-clade neutralizing antibodies (nAb) and T cell responses that were significantly more sustained compared to Bivalent Comirnaty (BC) mRNA vaccine booster. The persistence of RBD-specific follicular helper CD4^+^ T cells, germinal centre B cells, and long-lived plasma cells that facilitated affinity maturation in Clec9A^OMNI^-boosted mice, correlated with the detection of triple cross-reactive B cells that bind to ancestral SARS-CoV-2 ancestral, SARS-CoV-2 XBB.1.5 and SARS-CoV-1 RBD. Remarkably, intranasal boosting with Clec9A^OMNI^ generated robust and sustained mucosal immune responses in the upper and lower respiratory compartments, including RBD-specific IgA, cross-clade nAb and cellular immunity together with functional tissue-resident memory T cells, without compromising the systemic immune responses. Correspondingly, Clec9A^OMNI^ booster conferred superior protection against Omicron BA.1 compared to BC booster when challenge was performed at six months post-boost. Hence, Clec9A^OMNI^ is a promising nasal booster vaccine candidate that has the potential to mitigate pandemic threats from emerging sarbecoviruses.

**One Sentence Summary:** Nasal booster immunization with dendritic cell-targeting vaccine candidate in mRNA-vaccinated mice induced cross-clade, sustained, systemic and mucosal protective immunity.

## INTRODUCTION

Sarbecoviruses are a subgenus of *Betacoronavirus* and classified into four distinct evolutionary clades based on their receptor binding domain (RBD) sequence in spike protein: clade 1a [severe acute respiratory syndrome coronavirus (SARS-CoV-1) and related bat sarbecoviruses], clade 1b (SARS-CoV-2, related bat and pangolin sarbecoviruses), and clades 2 and 3 (bat sarbecoviruses) (*1*, *2*). Of concern, clade 1 members have been deemed of high zoonotic potential due to their ability to utilize human angiotensin-converting enzyme 2 (ACE2) as entry receptor (*2*). Hence, clade 1a and 1b sarbecoviruses pose a tangible pandemic threat.

The majority of the world population has developed varying levels of SARS-CoV-2 immunity through repeated infections and/or vaccinations. In particular, mRNA-based COVID-19 vaccines have been administered worldwide and found to induce a strong neutralizing antibody (nAb) response, which has been used as a correlate of protection against symptomatic COVID-19 (*3*, *4*). However, substantial waning of these antibodies has been observed, which rapidly reduces vaccine protective efficacy against re-infection (*5*). Additionally, given their intramuscular (i.m) route of administration, mRNA vaccines induce inadequate respiratory mucosal immunity, which is critical for protection against breakthrough infection and transmission (*6*, *7*). The highly inflammatory nature of lipid nanoparticles (LNP) that encapsulate the mRNA molecule, coupled with physical and chemical barriers within the respiratory tract, make current mRNA vaccine formulations unsuitable for intranasal (i.n) delivery (*8*, *9*). Lastly, the breadth of immune responses elicited by current mRNA COVID-19 vaccines, especially nAb, is limited towards clade 1b sarbecoviruses (*10*). Therefore, new-generation vaccines that can confer broad (cross-clade 1a and 1b), durable, systemic and mucosal protective immunity are imperative to prepare against future pandemics.

In recent years, dendritic cell (DC)-targeting strategies have been increasingly explored to develop novel vaccine candidates against infectious diseases, including SARS-CoV-2 and sarbecoviruses (*11–15*). Among which, targeting vaccine antigens to the C-type lectin-like receptor Clec9A expressed on conventional type 1 DCs (cDC1; mouse CD8α^+^ and human CD141^+^), has demonstrated great promise in pre-clinical animal models by inducing potent and durable immune responses upon a single-shot immunization, which have been associated with the generation of persistent antigen-specific follicular T-helper (T_FH_) cell, germinal center (GC), and antibody secreting cells (ASC) (*15–20*). Vaccine antigen candidates are genetically fused to the C-terminal end of each heavy chain of an anti-Clec9A monoclonal antibody (mAb). The superiority of Clec9A-targeting strategy compared to other DC-targeting approaches may be partly explained by the restricted expression of Clec9A on cDC1 subset, thereby resulting in longer circulation time of the antibody constructs and prolonged antigen presentation (*17*, *18*).

Previously, we engineered a Clec9A-RBD construct by fusing ancestral SARS-CoV-2 RBD to the heavy chains of anti-Clec9A mAb. We reported that single dose Clec9A-RBD systemic immunization elicited in mice potent and sustained immune responses against all SARS-CoV-2 variants, which translated into significant protection upon viral challenge (*15*).

Here, we evaluated Clec9A-RBD immunization as a booster approach to address the limitations faced by current mRNA vaccines. We evaluated the breadth and durability of the immune responses after booster immunization in mice that received two doses of Pfizer-BioNTech original mRNA vaccine (Comirnaty). Two Clec9A mAb constructs were generated, containing RBD from Omicron XBB.1.5 (Clec9A^XBB^) and SARS-CoV-1 (Clec9A^CoV1^) respectively. While boosting with Clec9A^XBB^ was expected to broaden the protective immune responses to Omicron variants, boosting with Clec9A^CoV1^ was expected to provide cross-clade (1a and 1b) protective immunity, based on previous studies reporting that SARS-CoV-1 survivors immunized with COVID-19 mRNA vaccines produced cross-clade nAb (*21*). Furthermore, CD103^+^ resident cDC1 expressing Clec9A are widely distributed throughout the respiratory tract which includes the lung and nasal mucosae (*22*, *23*). Hence, we explored for the first time the nasal delivery route of these Clec9A-RBD constructs to induce mucosal immune responses.

## RESULTS

### Boosting with Clec9A^XBB^ induced durable clade 1b-specific antibody responses while boosting with Clec9A^CoV1^ generated cross-clade antibody responses that waned over time

The breadth and durability of RBD-specific antibody responses were evaluated in mice that received two intramuscular (i.m) doses of Pfizer-BioNTech original Comirnaty mRNA vaccine three weeks apart, followed by systemic boosting three months later with Pfizer-BioNTech BA.4/5 bivalent Comirnaty (BC) mRNA vaccine (i.m), Clec9A^XBB^ or Clec9A^CoV1^ constructs; the Clec9A constructs were administered via the subcutaneous (s.c) route and adjuvanted with poly I:C **(Fig. 1A)**. Using a previously reported multiplex surrogate neutralizing assay (*21*), we observed that the three vaccine candidates effectively boosted the RBD-specific neutralizing antibody (nAb) titers **(Fig. 1B and S1)**. Furthermore, the nAb responses induced by Clec9A^XBB^ booster were sustained up to at least six months post-boost, while the responses induced in BC-boosted mice waned over time, consistent with previous reports (*24*) **(Fig. 1B and S1).** However, BC and Clec9A^XBB^ boosters both elicited poor cross-clade neutralization whereby significant nAb activity was restricted to clade 1b sarbecoviruses **(Fig. 1B and S1)**. In contrast, boosting with Clec9A^CoV1^ generated broad systemic nAb responses against clade 1a and 1b sarbecoviruses **(Fig. 1B and S1)**. However, the nAb responses elicited by C9A^CoV1^ booster were not as sustained as the Clec9A^XBB^ booster **(Fig. 1B and S1)**.

**Figure 1.**
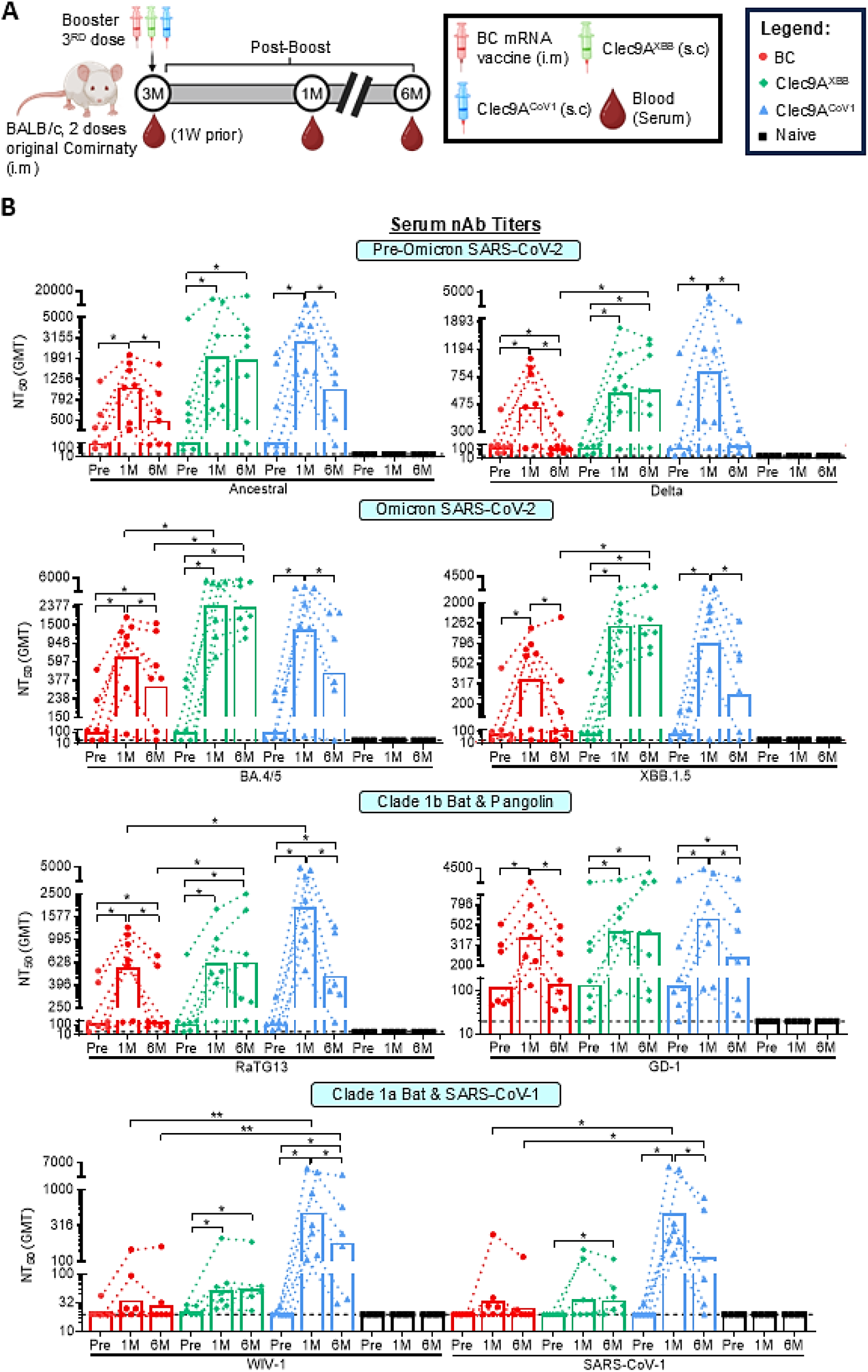
Breadth and durability of neutralizing antibody responses upon systemic booster immunization with mRNA vaccine (BC), Clec9A^XBB^ and Clec9A^CoV1^. (A) **Experimental design. Five to** six-week-old BALB/c mice (n = 6-7 per group) were immunized twice three weeks apart (0.05 µg per dose; i.m) with Pfizer-BioNTech original Comirnaty mRNA vaccine. Three months after the last immunization dose, mice were boosted either with Pfizer-BioNTech BA.4/5 bivalent Comirnaty (BC) mRNA vaccine (0.05 µg; i.m), Clec9A^XBB^ (10 µg adjuvanted with 50 µg poly I:C; s.c) or Clec9A^CoV1^ (10 µg adjuvanted with 50 µg poly I:C; s.c). A control group of non-immunized mice (naïve) was also included for baseline. At one-week before boost (Pre), and one– and six months post-boost (1M, 6M), blood was collected. **(B)** Serum nAb titers against eight sarbecoviruses from clades 1b and 1a were determined by multiplex sVNT. nAb titers were expressed as the reciprocal of the highest dilution that resulted in >50% inhibition (NT_50_). Symbols represent individual animals and data shown are geometric means. Statistical analysis: Non-parametric two-tailed Kruskal Walis test with Dunnett’s correction for multiple comparisons and Friedman test with Dunnett’s correction for multiple comparisons. *p < 0.05, **p < 0.01.

To further investigate the poorly sustained antibody responses induced by Clec9A^CoV1^, we monitored the durability of RBD-specific immune responses upon single shot immunization of naïve mice with Clec9A^XBB^ or Clec9A^CoV1^ **(Fig. S2A)**. Consistent with previous studies that have reported the relative weak humoral antigenicity of Omicron XBB variant compared to other variants (*25*, *26*), the RBD-specific IgG and nAb titers in Clec9A^XBB^-immunized mice were significantly lower than those measured in Clec9A^CoV1^-immunized animals. However, the antibody titers persisted for a longer duration in Clec9A^XBB^-immunized mice **(Fig. S2B&C)**. Interestingly, the cellular immune responses triggered upon restimulation of Clec9A^CoV1^-immunized splenocytes were much stronger compared to the Clec9A^XBB^-immunized group, which were associated with robust CD4^+^ type 1 cytokine responses **(Fig. S2D)**. Despite this strong cellular response, the RBD-specific T_FH_ and GC B cell responses in the spleen from Clec9A^CoV1^-immunized mice were barely detectable at six months post-immunization **(Fig. S2E&F)**. In addition, in Clec9A^CoV1^-immunized mice, RBD-specific antibody secreting cells (ASC) in the bone marrow (BM) were skewed towards the non-long-lived plasma cell (non-LLPC) subset **(Fig. S2G)**. In contrast, Clec9A^XBB^-immunized mice displayed a significantly greater number of RBD-specific T_FH_ and GC B cells in their spleen at six months post-immunization, coupled with increased differentiation of antigen-specific ASC into LLPC in the BM **(Fig. S2E-G)**. These observations provide a likely explanation for the differential persistence of the antibody responses observed between Clec9A^CoV1^ and Clec9A^XBB^-immunized mice.

Together, these results showed that Clec9A^XBB^ booster in mRNA-immunized mice elicited a sustained humoral response that was limited in breadth, while Clec9A^CoV1^ booster produced a cross-clade nAb response that did not persist as long as the Clec9A^XBB^ booster, likely due to poor intrinsic ability to induce persistent T_FH_ and GC B cell responses. Hence, these observations suggested that the amino acid make-up of RBD antigen influenced the immune responses.

### Broad and durable RBD-specific humoral responses and protection after systemic booster immunization with Clec9A^OMNI^

To mitigate the respective limitations faced by Clec9A^XBB^ and Clec9A^CoV1^, we co-administered both constructs, at 1:1 or 4:1 ratio (Clec9A^XBB^: Clec9A^CoV1^) **(Fig. S3A)**. Results indicated that while both ratio formulations induced comparable nAb titers against both sarbecovirus clades at 3-week post-boost **(Fig. S3B)**, the 4:1 ratio formulation elicited more robust cross-clade T cell responses, although the differences did not reach statistical significance **(Fig. S3C)**.

Using the 4:1 dose combination (hereby referred to as Clec9A^OMNI^), we carried out in-depth characterization of the breadth and durability of the humoral immune responses induced upon s.c boosting **(Fig. 2A)**. Results indicated that Clec9A^OMNI^ booster induced systemic cross-clade nAb responses **(Fig. 2B and S4)**, along with the presence in the spleen of RBD-specific switched immunoglobulin positive (swIg^+^) B cells that were cross-reactive towards ancestral SARS-CoV-2, XBB.1.5 and SARS-CoV-1 RBD **(Fig. 2C)**. On the contrary, the cross-reactivity of antigen-specific swIg^+^ B cells in BC-boosted animals were limited to ancestral SARS-CoV-2 and XBB.1.5 RBD only **(Fig. 2C)**, consistent with the limited neutralizing activity observed against clade 1a sarbecoviruses **(Fig. 2B and S4)**. Additionally, the cross-clade systemic nAb responses induced by Clec9A^OMNI^ booster were sustained at six months post-boost **(Fig. 2B)**, which correlated with greater number of RBD-specific T_FH_ and GC B cells in the spleen **(Fig. 2D)**, and greater number of RBD-specific LLPC in the BM **(Fig. 2E)** at six months post-boost compared to BC-boosted animals.

**Figure 2.**
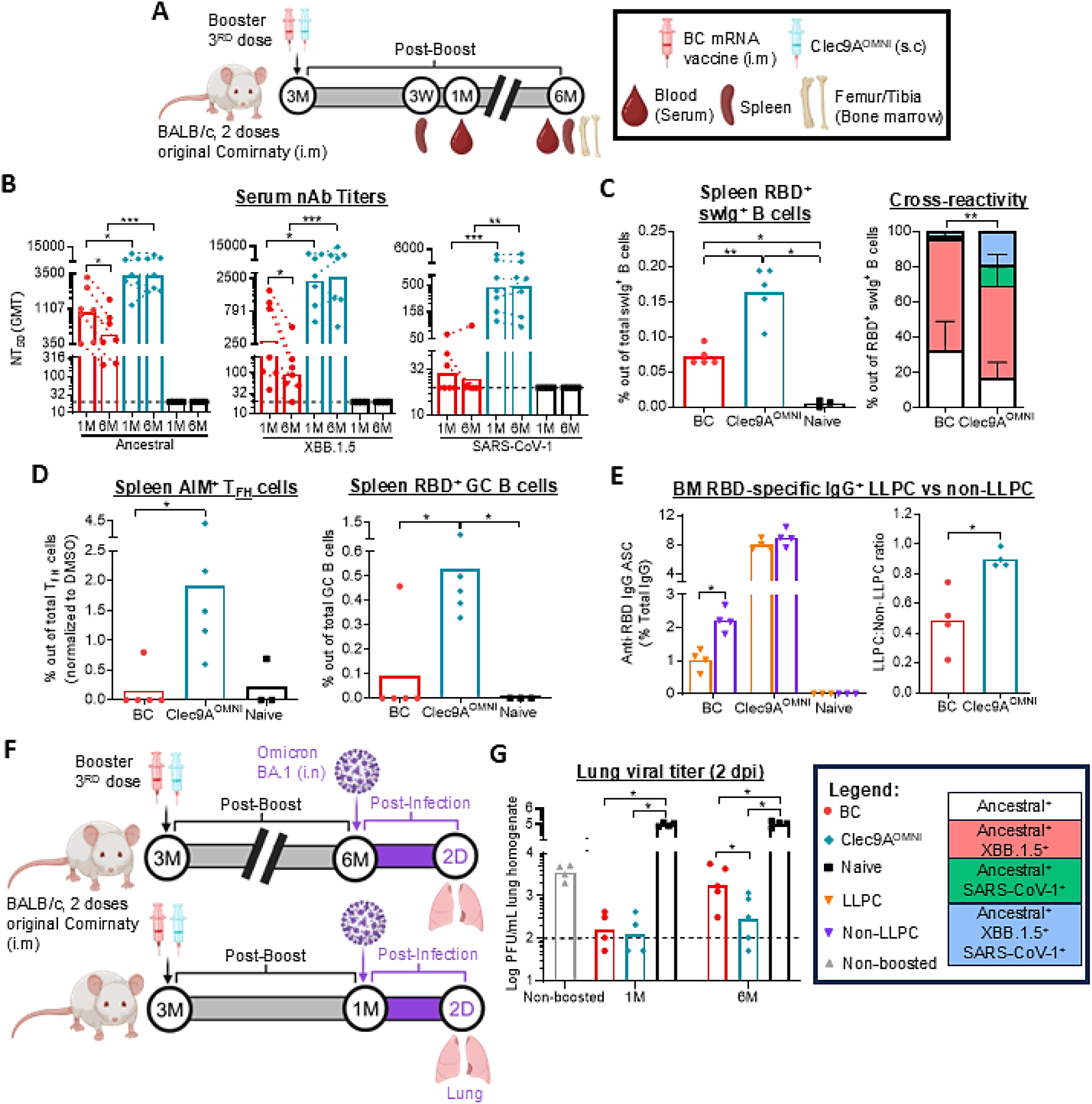
Breadth and durability of RBD-specific humoral responses and protection upon systemic booster immunization with Clec9A^OMNI^ versus BC mRNA vaccine. **(A)** Immunogenicity experiment design. Five to six-week-old BALB/c mice (n = 4-7 per group) were immunized twice three weeks apart (0.05 µg per dose; i.m) with original Comirnaty mRNA vaccine. Three months after the second immunization dose, mice were boosted with either BC mRNA vaccine (0.05 µg; i.m), or Clec9A^OMNI^ (8 µg Clec9A^XBB^ + 2 µg Clec9A^CoV1^ adjuvanted with 50 µg poly I:C; s.c). **(B)** Serum nAb titers (NT_50_) against ancestral SARS-CoV-2, XBB.1.5 and SARS-CoV-1 at one– and six months post-boost were determined by multiplex sVNT. **(C)** Percentage of ancestral SARS-CoV-2 RBD^+^ swIg^+^ B cells in spleen at three weeks post-boost were determined by flow cytometry. Cross-reactivity of these antigen-specific BC9A cells towards XBB.1.5 and SARS-CoV-1 RBD were also shown. **(D)** Percentages of activation-induced markers positive (AIM^+^) T_FH_ (CD4^+^ CXCR5^+^ PD-1^+^ CD69^+^ CD154^+^) and RBD^+^ GC B cells (B220^+^ IgD^-^ GL-7^+^ CD95^+^ RBD^++^) in spleen at six months post-boost were determined by flow cytometry. **(E)** Frequency of BM RBD-specific IgG^+^ LLPC and non-LLPC (normalized to total IgG) at six months post-boost was determined by B cell ELISPOT. SFU = spot forming unit. **(F)** Challenge experiment design. At one– and six months post-boost, mice were challenged with 10^6^ PFU Omicron BA.1 via the i.n route. At two days post-infection (dpi), mice were euthanized, and lungs were harvested and homogenized. **(G)** Lung viral titers at two dpi were quantified via plaque assay. The dashed line represents the LOD at 2 log_10_, and samples below the LOD were given an arbitrary value corresponding to half the LOD value at 1.7 log_10_. **(B-E, G)** Symbols represent individual animals and data shown are **(B)** geometric means and **(C-E, G)** means ± **(C)** standard deviation (SD). Statistical analysis: Non-parametric two-tailed **(B, C, E)** Mann-Whitney test, **(B)** Wilcoxon matched-pairs signed rank test, and **(C, D, G)** Kruskal Walis test with Dunnett’s correction for multiple comparisons. *p < 0.05, **p < 0.01, ***p < 0.001.

The protective efficacy of Clec9A^OMNI^ systemic booster was also investigated. Mice were challenged nasally with Omicron BA.1 virus at either one-or six months post-boost, and the lung viral titers were measured **(Fig. 2F)**. Compared to non-boosted animals, mice boosted with either BC or Clec9A^OMNI^ had comparable and significantly reduced lung viral titers upon challenge performed at one-month post-boost **(Fig. 2G)**. However, when the challenge was performed at six months post-boost, the lung viral titers were significantly lower in Clec9A^OMNI^-boosted mice compared with BC-boosted mice **(Fig. 2G)**.

Together, these data indicated that sc. booster immunization with Clec9A^OMNI^ induced cross-clade and durable nAb responses that conferred superior long-term protection compared to BC mRNA booster.

### Nasal Clec9A^OMNI^ booster immunization generated robust, cross-clade RBD-specific systemic and mucosal humoral responses

The ability to induce protective immunity at the respiratory mucosa represents a highly desired and notable advancement over current systemic vaccination methods for providing strong protection and reducing transmission between individuals. Thus, we investigated the suitability of the nasal route of Clec9A^OMNI^ booster immunization **(Fig. 3A)**. At one-month post-boost, both BC (i.m) and Clec9A^OMNI^ (i.n) boosters triggered strong serum anti-RBD IgG responses, although Clec9A^OMNI^ induced greater binding antibody titers against SARS-CoV-1 RBD **(Fig. 3B)**. Consistently, both boosters produced RBD-specific IgG^+^ antibody secreting cells (ASC) in the spleen and BM, where ASC in BC-boosted mice were mainly reactive to ancestral SARS-CoV-2 and XBB.1.5 RBD, while Clec9A^OMNI^ elicited higher frequency of ASC that were reactive to ancestral SARS-CoV-2, XBB.1.5 and SARS-CoV-1 RBD **(Fig. 3C)**.

**Figure 3.**
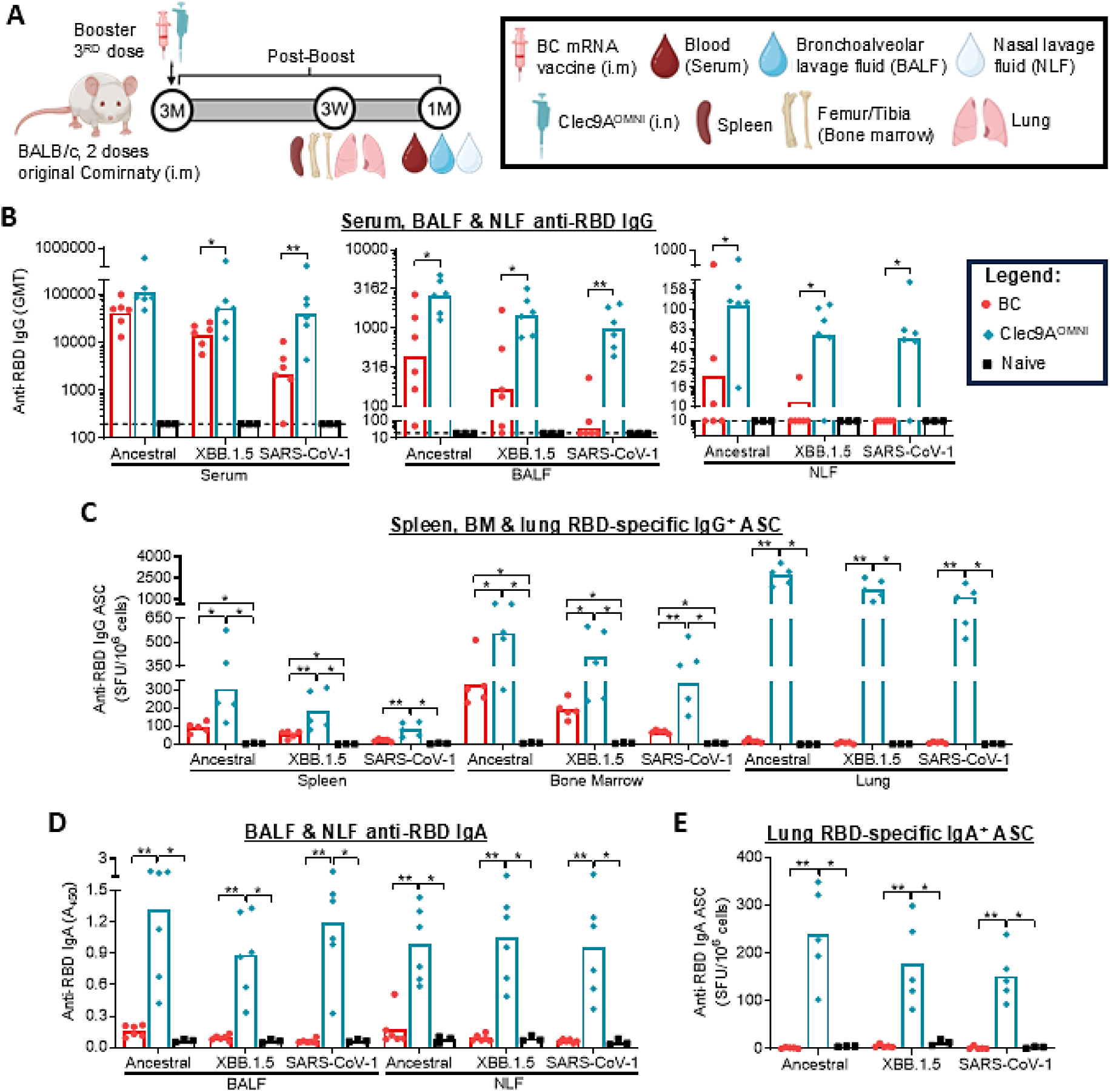
RBD-specific IgG and IgA responses upon nasal booster immunization with Clec9A^OMNI^ versus systemic booster immunization with BC mRNA vaccine. **(A)** Immunogenicity experiment design. Five to six-week-old BALB/c mice (n = 5-6 per group) were immunized twice three weeks apart with original Comirnaty mRNA vaccine (0.05 µg per dose; i.m). Three months after the second immunization dose, mice were boosted with either BC mRNA vaccine (0.05 µg; i.m), or Clec9A^OMNI^ (4 µg Clec9A^XBB^ + 1 µg Clec9A^CoV1^ adjuvanted with 50 µg poly I:C; i.n). **(B)** Serum, BALF and NLF anti-RBD IgG titers against ancestral SARS-CoV-2, XBB.1.5 and SARS-CoV-1 RBD at one-month post-boost were determined by ELISA. The dashed line represents the LOD at 200, 20 and 10 respectively. **(C)** Frequency of spleen, BM and lung RBD-specific IgG^+^ ASC reactive to ancestral SARS-CoV-2, XBB.1.5 and SARS-CoV-1 RBD at three weeks post-boost was determined by B cell ELISPOT. SFU = spot forming unit. **(D)** BALF and NLF anti-RBD IgA titers against ancestral SARS-CoV-2, XBB.1.5 and SARS-CoV-1 RBD at one-month post-boost were determined by ELISA. **(E)** Frequency of lung RBD-specific IgA^+^ ASC reactive to ancestral SARS-CoV-2, XBB.1.5 and SARS-CoV-1 RBD at three weeks post-boost was determined by B cell ELISPOT. **(B-E)** Symbols represent individual animals and data shown are **(B)** geometric means and **(C-E)** means. Statistical analysis: Non-parametric two-tailed **(B)** Mann-Whitney test, and **(C-E)** Kruskal Walis test with Dunnett’s correction for multiple comparisons. *p < 0.05, **p < 0.01.

Low levels of anti-RBD IgG were detected in bronchoalveolar and nasal lavage fluids (BALF, NLF) from BC-boosted mice **(Fig. 3B)**, with undetectable RBD-specific IgG^+^ ASC in the lung tissues **(Fig. 3C)**, suggesting that the IgG detected in the lavage fluids were likely spillover from the circulation as previously proposed (*7*, *9*, *27*). Clec9A^OMNI^ booster instead generated significantly higher anti-RBD IgG titers against ancestral SARS-CoV-2, XBB.1.5 and SARS-CoV-1 RBD in the BALF and NLF **(Fig. 3B)**, with the detection of triple cross-reactive RBD-specific IgG^+^ ASC in lung tissues, supporting successful priming of antigen-specific B cells and antibody responses in the respiratory mucosa **(Fig. 3C)**. Furthermore, while undetectable following BC booster, nasal boosting with Clec9A^OMNI^ induced anti-RBD IgA in BALF and NLF, coupled with RBD-specific IgA^+^ ASC in the lung tissues **(Fig. 3D&E)**.

The systemic and respiratory nAb responses were also measured at one-month post-boost **(Fig. 4A)**. Although i.m BC booster induced potent systemic nAb responses against clade 1b sarbecoviruses, the BC-boosted responses were very limited in the BALF and NLF **(Fig. 4B-D and S5A-C)**, consistent with undetectable levels of RBD-specific swIg^+^ B cells in the lung and NALT tissues **(Fig. 4E)**. In contrast, i.n Clec9A^OMNI^ booster elicited robust and cross-clade serum, BALF and NLF nAb responses **(Fig. 4B-D and S5A-C)**, that correlated well with the generation of cross-reactive RBD-specific swIg^+^ B cells to ancestral SARS-CoV-2, XBB.1.5 and SARS-CoV-1 RBD in the spleen, lung and NALT tissues **(Fig. 4E)**.

**Figure 4.**
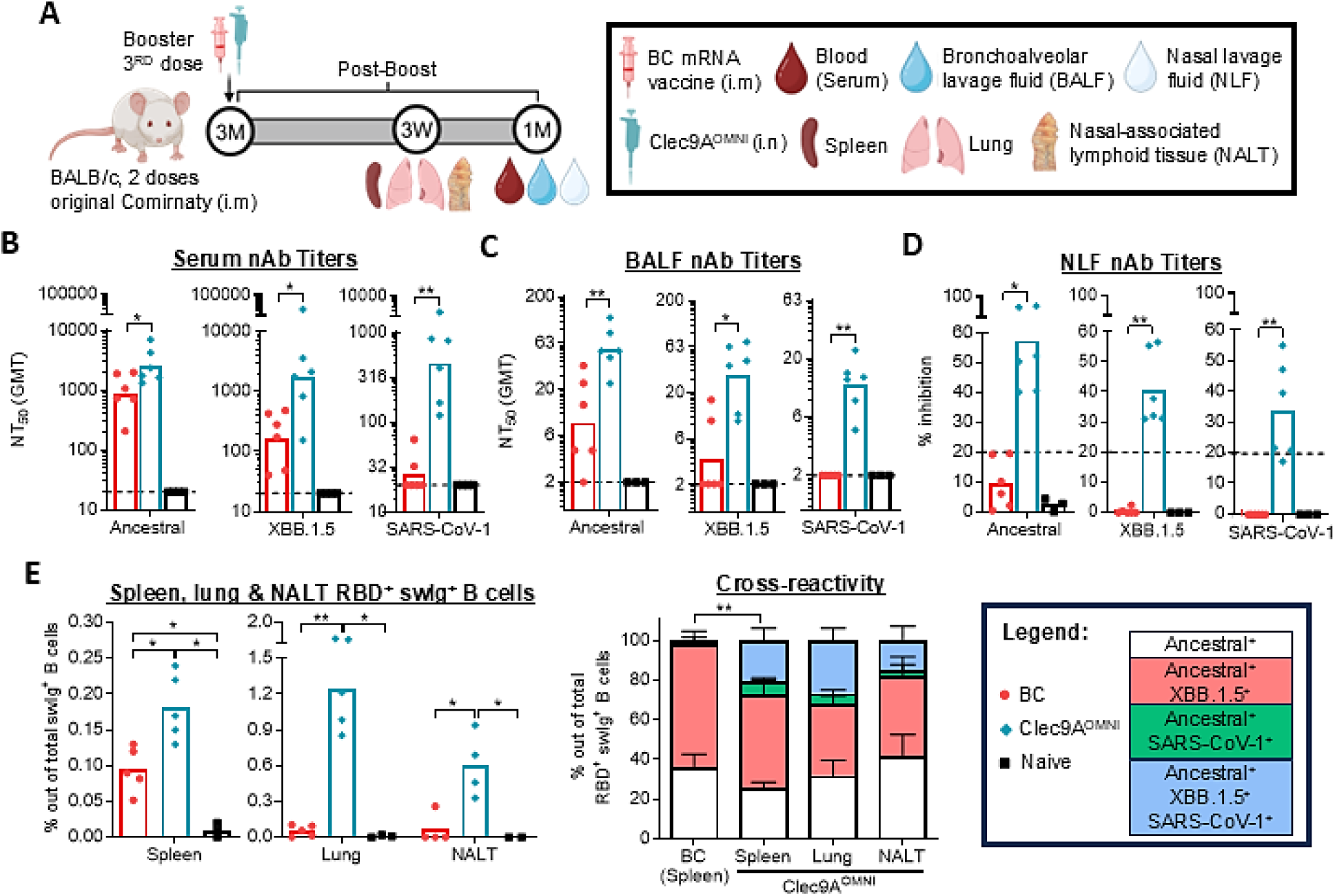
Breadth of neutralizing antibody responses upon nasal booster immunization with Clec9A^OMNI^ versus systemic booster immunization with BC mRNA vaccine. **(A)** Immunogenicity experiment design. Five to six-week-old BALB/c mice (n = 4-6 per group) were immunized twice three weeks apart with original Comirnaty mRNA vaccine (0.05 µg per dose; i.m). Three months after the second immunization dose, mice were boosted with either BC mRNA vaccine (0.05 µg; i.m), or Clec9A^OMNI^ (4 µg Clec9A^XBB^ + 1 µg Clec9A^CoV1^ adjuvanted with 50 µg poly I:C; i.n). **(B-D)** At one-month post-boost, mice were euthanized and blood, BALF and NLF were collected. Serum **(B)**, BALF **(C)** and NLF **(D)** nAb titers (serum, BALF: NT_50_, NLF: % inhibition at 1:2 dilution) against ancestral SARS-CoV-2, XBB.1.5 and SARS-CoV-1 were determined by multiplex sVNT. **(E)** Mice were euthanized at three weeks post-boost, and spleen, lung and NALT tissues were harvested. The percentage of ancestral SARS-CoV-2 RBD^+^ swIg^+^ B cells was determined by flow cytometry. Cross-reactivity of these antigen-specific B cells towards XBB.1.5 and SARS-CoV-1 RBD were shown. (**B-E)** Symbols represent individual animals and data shown are **(B, C)** geometric means and **(D, E)** means ± SD. Statistical analysis: Non-parametric two-tailed **(B-E)** Mann-Whitney test and **(E)** Kruskal Walis test with Dunnett’s correction for multiple comparisons. *p < 0.05, **p < 0.01.

Together, the results demonstrated that i.n booster immunization with Clec9A^OMNI^ produced potent and broad RBD-specific systemic and mucosal humoral responses against clade 1a and 1b sarbecoviruses, compared to i.m BC mRNA booster which induced limited respiratory humoral immunity.

### Nasal Clec9A^OMNI^ booster immunization induced cross-clade RBD-specific systemic and mucosal polyfunctional cellular responses

The antigen-specific systemic and mucosal T cell responses were next characterized two weeks after i.n boosting with Clec9A^OMNI^, relative to BC i.m booster **(Fig. 5A)**. Although both boosters elicited detectable RBD-specific systemic cellular responses upon restimulation of splenocytes with sarbecovirus RBD peptides from both clades 1a and 1b, these responses were much stronger and more polyfunctional in Clec9A^OMNI^-boosted mice compared to BC-boosted mice **(Fig. 5B)**. Furthermore, while undetectable in the BC boosted group, mice boosted with i.n Clec9A^OMNI^ produced cross-clade RBD-specific mucosal mono– and polyfunctional T cell responses upon restimulation of lung and NALT cells with sarbecovirus RBD peptides **(Fig. 5C and 5D)**.

**Figure 5.**
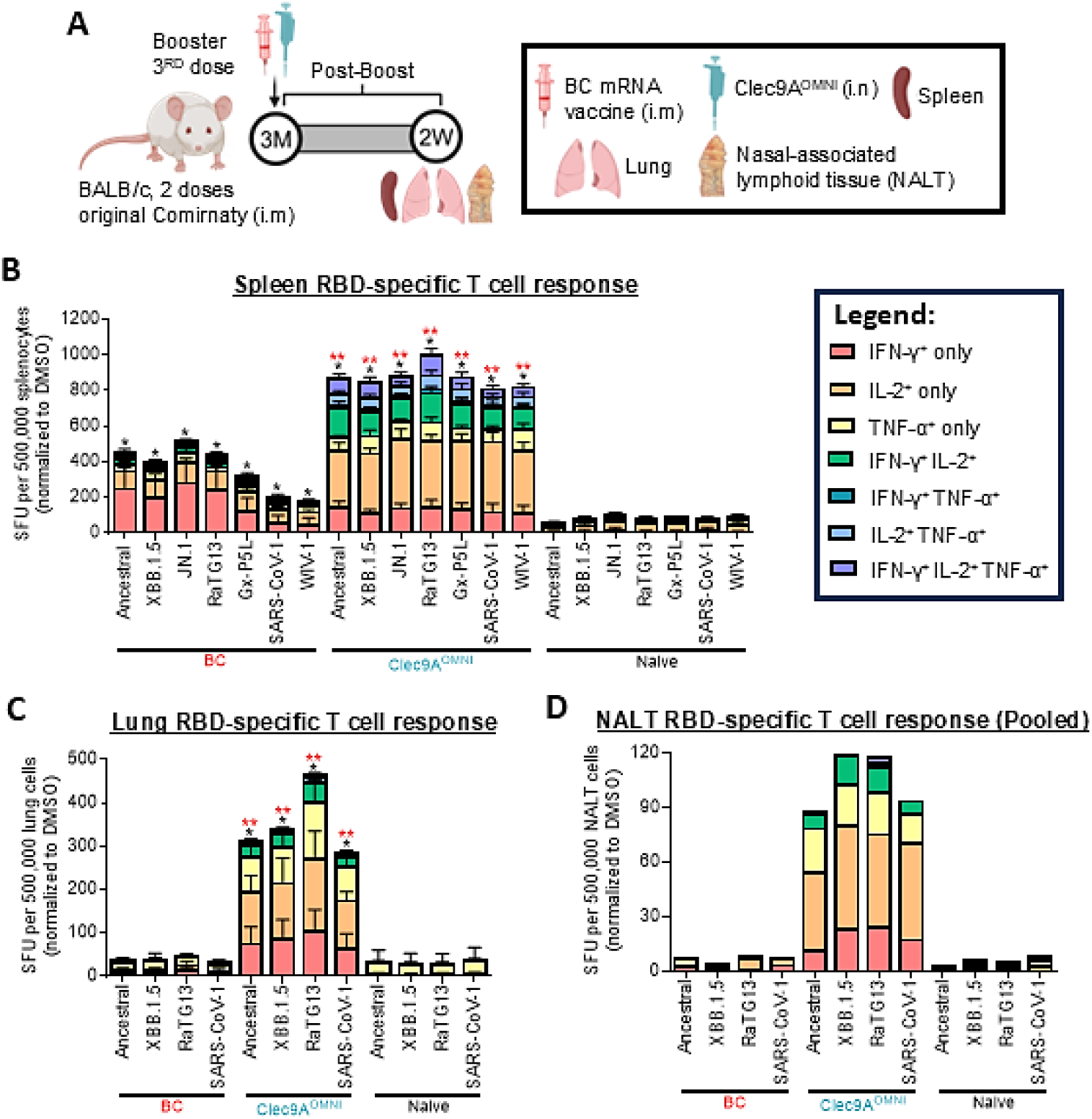
RBD-specific cellular responses upon nasal booster immunization with Clec9A^OMNI^ versus systemic booster immunization with BC mRNA vaccine. **(A)** Immunogenicity experiment design. Five to six-week-old BALB/c mice (n = 5 per group) were immunized twice three weeks apart with original Comirnaty mRNA vaccine (0.05 µg per dose; i.m). Three months after the second immunization dose, mice were boosted with either BC mRNA vaccine (0.05 µg; i.m), or Clec9A^OMNI^ (4 µg Clec9A^XBB^ + 1 µg Clec9A^CoV1^ adjuvanted with 50 µg poly I:C; i.n). At two weeks post-boost, mice were euthanized, and spleen, lung and NALT tissues were harvested. **(B-D)** Frequency of IFN-γ, IL-2 and/or TNF-α secreting **(B)** splenocytes, **(C)** lung cells, and **(D)** pooled NALT cells was determined by FluoroSPOT upon restimulation with ancestral SARS-CoV-2, XBB.1.5, JN.1, RaTG13, Gx-P5L, SARS-CoV-1 and WIV-1 RBD peptides. SFU = spot forming units. Data shown are **(B-D)** means ± **(B, C)** SD. Statistical analysis: Non-parametric two-tailed Kruskal Walis test with Dunnett’s correction for multiple comparisons. *p < 0.05, **p < 0.01.

Together, these findings showed that i.n boosting with Clec9A^OMNI^ elicited robust and cross-clade RBD-specific systemic and mucosal (upper and lower respiratory tract) polyfunctional T cell responses against sarbecoviruses.

### Nasal Clec9A^OMNI^ booster immunization induced durable, cross-clade, systemic and mucosal immune responses that were protective

The immunogenicity and protective efficacy of Clec9A^OMNI^ nasal booster were next evaluated at six months post-boost **(Fig. 6A)**. Potent IgG responses specific to ancestral SARS-CoV-2, XBB.1.5 and SARS-CoV-1 RBD were detected in the serum, BALF and NLF from i.n Clec9A^OMNI^-boosted animals, with titers that were significantly higher than those measured in i.m BC-boosted animals **(Fig. 6B)**. Of note, the difference in antibody titers between both groups were greater than at one-month post-boost **(Fig. 3B and 6B)**, due to the waning effect observed in BC-boosted mice. Moreover, while still undetectable in BC-boosted mice, cross-clade anti-RBD IgA responses persisted in the BALF and NLF from Clec9A^OMNI^-boosted mice at six months post-boost **(Fig. 6C)**.

**Figure 6.**
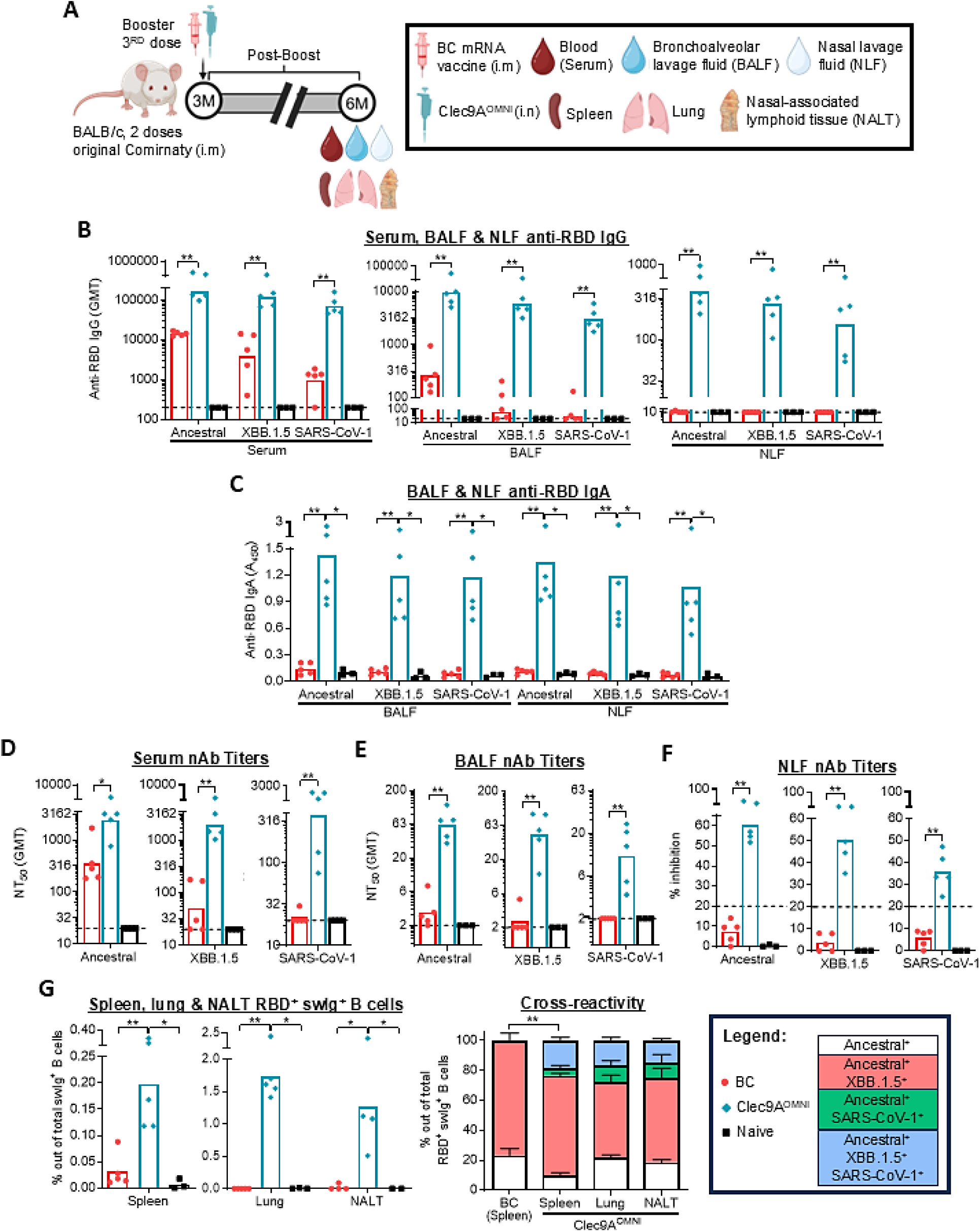
Durability of RBD-specific antibody responses upon nasal booster immunization with Clec9A^OMNI^ versus systemic booster immunization with BC mRNA vaccine. **(A)** Immunogenicity experiment design. Five to six-week-old BALB/c mice (n = 6 per group) were immunized twice three weeks apart with original Comirnaty mRNA vaccine (0.05 µg per dose; i.m). Three months after the second immunization dose, mice were boosted with either BC mRNA vaccine (0.05 µg; i.m), or Clec9A^OMNI^ (4 µg Clec9A^XBB^ + 1 µg Clec9A^CoV1^ adjuvanted with 50 µg poly I:C; i.n). Mice were bled and euthanized at six months post-boost. **(B)** Serum, BALF and NLF anti-RBD IgG titers against ancestral SARS-CoV-2, XBB.1.5 and SARS-CoV-1 RBD were determined by ELISA. The dashed line represents the LOD at 200, 20 and 10 respectively. **(C)** BALF and NLF anti-RBD IgA titers against ancestral SARS-CoV-2, XBB.1.5 and SARS-CoV-1 RBD were determined by ELISA. **(D)** Serum, **(E)** BALF and **(F)** NLF nAb titers (serum, BALF: NT_50_, NLF: % inhibition at 1:2 dilution) against ancestral SARS-CoV-2, XBB.1.5 and SARS-CoV-1 were determined by multiplex sVNT. **(G)** Percentage of ancestral SARS-CoV-2 RBD^+^ swIg^+^ B cells in spleen, lung and NALT tissues was determined by flow cytometry. Cross-reactivity of these antigen-specific B cells towards XBB.1.5 and SARS-CoV-1 RBD were also shown. (**B-G)** Symbols represent individual animals and data shown are **(B, D, E)** geometric means and **(C, F, G)** means. Statistical analysis: Non-parametric two-tailed **(B, D-G)** Mann-Whitney test, and **(C, G)** Kruskal Walis test with Dunnett’s correction for multiple comparisons. *p < 0.05, **p < 0.01.

Similarly, robust systemic and mucosal cross-clade nAb responses were still detected in the serum, BALF and NLF of Clec9A^OMNI^-boosted mice at six months post-boost **(Fig. 6D-F and S6A-C)**, correlating with the persistence of cross-reactive RBD-specific swIg^+^ B cells towards ancestral SARS-CoV-2, XBB.1.5 and SARS-CoV-1 RBD in spleen, lung and NALT tissues **(Fig. 6G)**. In contrast, BC-boosted mice displayed significantly lower potency and breadth in serum nAb responses, coupled with largely undetectable neutralizing activities in the BALF and NLF **(Fig. 6D-F and S6A-C)**. These observations were consistent with the limited presence of cross-reactive RBD-specific swIg^+^ B cells in all three immune compartments of BC-boosted mice **(Fig. 6G)**.

The durable systemic and mucosal antibody responses following i.n boosting with Clec9A^OMNI^ were associated with the presence of RBD-specific T_FH_ and GC B cells in the spleen and lung tissues at six months post-boost, coupled with high number of RBD-specific IgG^+^ LLPC in the BM **(Fig. 7A-C)**. Moreover, CD4^+^ and CD8^+^ T cells with functional tissue-resident memory (T_RM_) phenotype were detected in the lung and NALT tissues harvested at three-month post-boost **(Fig. 7D)**.

**Figure 7.**
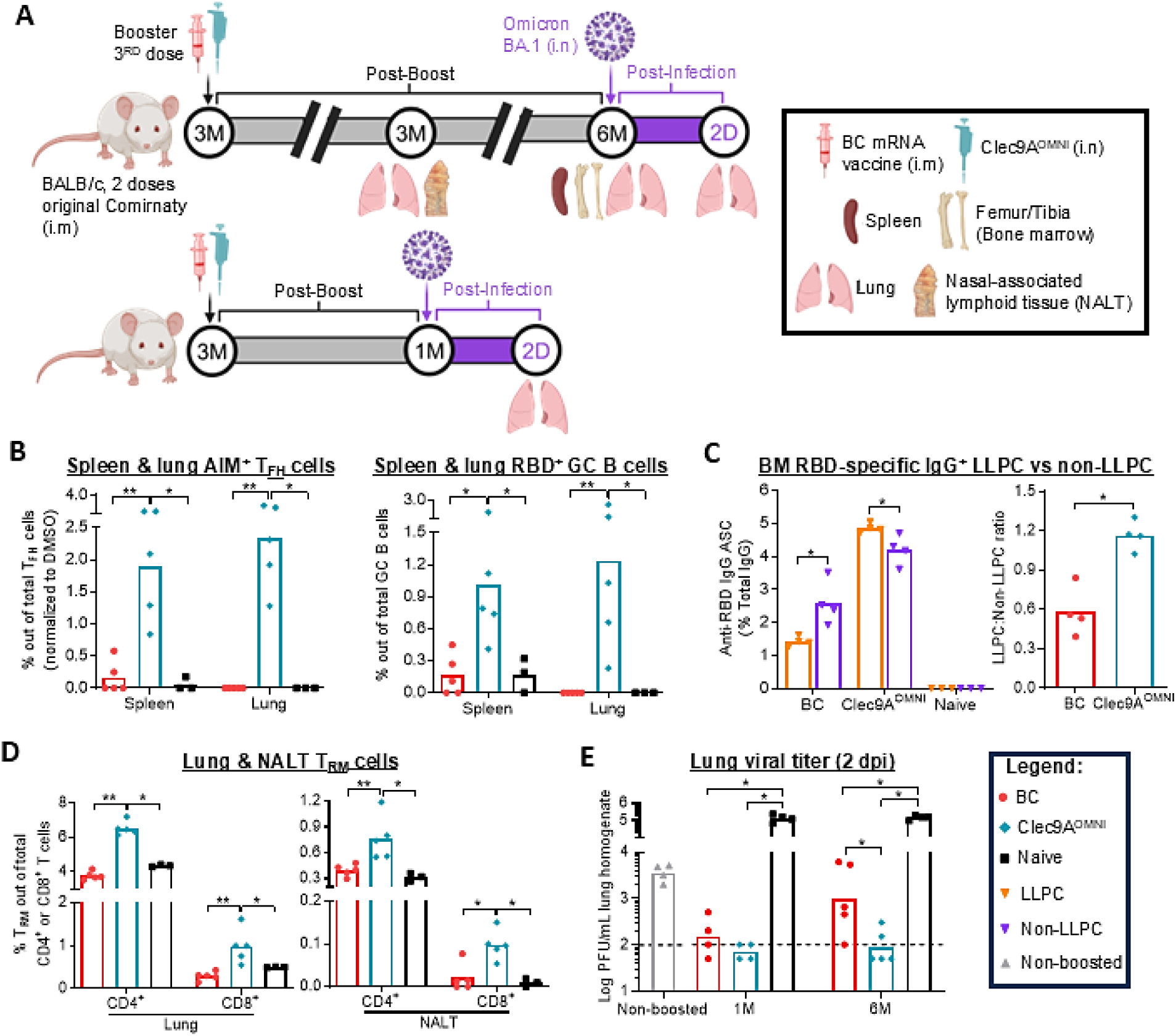
Durability of RBD-specific immune responses and protection upon nasal booster immunization with Clec9A^OMNI^ versus systemic booster immunization with BC mRNA vaccine. **(A)** Immunogenicity and challenge experiment design. Five to six-week-old BALB/c mice (n = 4-5 per group) were immunized twice three weeks apart with original Comirnaty mRNA vaccine (0.05 µg per dose; i.m). Three months after the second immunization dose, mice were boosted with either BC mRNA vaccine (0.05 µg; i.m), or Clec9A^OMNI^ (4 µg Clec9A^XBB^ + 1 µg Clec9A^CoV1^ adjuvanted with 50 µg poly I:C; i.n). **(B)** Percentages of AIM^+^ T_FH_ and RBD^+^ GC B cells in spleen and lung tissues at six months post-boost were determined by flow cytometry. **(C)** Frequency of BM RBD-specific IgG^+^ LLPC and non-LLPC (normalized to total IgG) at six months post-boost was determined by B cell ELISPOT. SFU = spot forming unit. **(D)** Percentage of lung and NALT CD4^+^ (CD69^+^ CD49a^+^) and CD8^+^ (CD69^+^ CD103^+^ CD49a^+^) T_RM_ cells at three months post-boost was determined by flow cytometry. **(E)** Mice were challenged at one-or six months post-boost with 10^6^ PFU Omicron BA.1 via the i.n route. At two dpi, mice were euthanized, and lungs were harvested and homogenized. Lung viral titers at two dpi were quantified via plaque assay. The dashed line represents the LOD at 2 log_10_, and samples below the LOD were given an arbitrary value corresponding to half the LOD value at 1.7 log_10_. **(B-E)** Symbols represent individual animals and data shown are means. Statistical analysis: Non-parametric two-tailed **(C)** Mann-Whitney test and **(B, D, E)** Kruskal Walis test with Dunnett’s correction for multiple comparisons. *p < 0.05, **p < 0.01.

The persistent systemic and mucosal humoral and cellular immunity measured in Clec9A^OMNI^-boosted (i.n) animals translated into strong protection against BA.1 challenge, whereby the lung viral titers remained at the detection limit even when challenge was performed at six months post-boost **(Fig. 7E)**. Conversely, BC booster provided reduced long-term protective efficacy against BA.1 infection **(Fig. 7E)**, which was associated with limited induction and persistence of cellular subsets, and antigen-specific ASC skewed towards non-LLPC **(Fig. 7B-D).**

Collectively, these results indicated that boosting nasally with Clec9A^OMNI^ induced cross-clade RBD-specific humoral and T cell responses in both systemic and mucosal (respiratory) compartments. Furthermore, immune responses remained highly durable with the persistence of antibodies (IgG, IgA and nAb), long-lived RBD-specific B cell subsets, and the establishment of respiratory mucosa-resident cellular memory. These immune responses conferred sustained and robust protection against viral challenge.

## DISCUSSION

Given that majority of the global population has developed some level of immunity to SARS-CoV-2, either through vaccination, infection, or both, the next generation of vaccine candidates targeting sarbecoviruses should broaden existing SARS-CoV-2 immunity and induce long-lasting responses. In this study, we showed that boosting simultaneously with Omicron XBB.1.5 and SARS-CoV-1 RBD using a Clec9A-targeting mAb (Clec9A^OMNI^) broadened the immune responses to clade 1a and 1b sarbecoviruses in mRNA Comirnaty-vaccinated mice. In Clec9A^OMNI^-boosted animals, we detected a subset of B cells in the spleen and respiratory mucosa that were triple cross-reactive to RBD from ancestral SARS-CoV-2, XBB.1.5 and SARS-CoV-1, which strongly supported the production of broadly neutralizing antibodies (nAb), in addition to the production of clade-specific nAb. Triple cross-reactive B cells were not detected in mice boosted with BC mRNA vaccine. Our finding supported that antigenic exposure to both sarbecovirus clades is required to elicit cross-clade humoral breadth, which is consistent with the low degree of conservation between RBD sequences (*28*), and despite the presence of conserved B cell epitopes between clade 1a and 1b sarbecovirus (*29*, *30*). Although COVID-19 mRNA vaccines were shown to induce RBD-specific B cell clones that produced cross-clade nAb targeting these conserved epitopes (*30–32*), these B cell clones are extremely rare. Hence, boosting with Omicron-based mRNA vaccine is likely to limit the expansion of these cross-clade B cell clones, and instead favors the recall of B cells that recognize epitopes that are shared within the clade 1b and that are in greater abundance (*29*, *30*). To avoid this immune imprinting phenomenon, it was proposed that boosting with SARS-CoV-1 RBD increases the selection stringency towards those rare cross-clade B cell clones and avoids the recall of clade 1b-specific B cell clones (*33*). Consistently, previous studies have reported the induction of pan-sarbecovirus nAb responses through sequential cross-clade vaccination strategies (*34*, *35*). Likewise, SARS-CoV-2 infection or vaccination of individuals with pre-existing SARS-CoV-1 immunity triggered cross-clade nAb responses (*21*).

While boosting with Clec9A^CoV1^ alone was sufficient to induce cross-clade humoral responses in Comirnaty mRNA-vaccinated mice, the RBD-specific T cell responses were mostly specific to SARS-CoV-1. On the other hand, boosting with Clec9A^XBB^ alone generated robust cellular responses against SARS-CoV-2 variants but weak T cell responses against SARS-CoV-1. Cross-clade T cell responses were only produced upon simultaneous boosting with the two Clec9A constructs (Clec9A^OMNI^). This finding suggested low T cell cross-reactivity between both sarbecovirus clades and indicated that the breadth of cellular immune responses induced by Clec9A^OMNI^ is likely mediated by a combination of *de novo* responses against clade 1a sarbecoviruses, coupled with recall of cross-reactive T cells from prior SARS-CoV-2 mRNA vaccination against clade 1b sarbecoviruses. Indeed, studies have reported low amino acid conservation between the immunodominant spike peptide pools of SARS-CoV-2 and SARS-CoV-1 (*36*). Therefore, to elicit both cross-clade nAb and T cell responses following prior mRNA Comirnaty vaccination, formulations that contain both clade 1a and 1b sarbecovirus RBD (*i.e* chimeric antigen, mosaic nanoparticle, and multivalent cocktail vaccines) may represent better booster strategies (*34*, *37–39*).

One of the unique features of the Clec9A-targeting strategy is the ability to induce exceptionally sustained immune responses, which have been associated with the persistence of GC B cells and follicular helper T cells (T_FH_) (*15*, *16*). Consistently, Clec9A^OMNI^ booster induced sustained antibody responses, which correlated with the persistence of RBD-specific T_FH_ cells, GC B cells and increased differentiation into LLPC. In contrast, boosting with Comirnaty bivalent mRNA vaccine failed to establish LLPC in the bone marrow, where the majority of RBD-specific ASC displayed a non-LLPC phenotype, and in line with a previous report (*40*). The differentiation of GC B cells into LLPC is dependent on T_FH_ cell-derived cytokines, costimulatory and regulatory signals (*41*, *42*). We found that immunization with Clec9A^XBB^ induced a strong and persistent T_FH_ response, which was associated with sustained antibody titers. While immunization with Clec9A^CoV1^ induced a less potent and durable T_FH_ cell response, which was associated with less persistent antibody titers. This observation suggested that the RBD antigenic sequence fused to the anti-Cle9A antibody influenced the immune responses. While the exact mechanism remains to be determined, variation in spike-specific T_FH_ responses have been reported between closely related SARS-CoV-2 variants (*43*). Clec9A^CoV1^ immunization triggered a strong CD4^+^ T_H_1 but poor T_FH_ response. Whereas Clec9A^XBB^ immunization elicited a more balanced repertoire of T_H_1 and T_FH_ responses. Cytokines such as Type-I IFN, IL-6 and IL-12 drive the CD4^+^ T cell polarization towards a T_FH_-biased phenotype (*41*, *44*). Additionally, T cell receptor signal strength can also influence the polarization of CD4^+^ T-helper subsets where strong signaling was shown to promote terminal T_H_1 differentiation over T_FH_ (*45*, *46*). Hence, the avidity of T cell interaction with their respective MHC-peptide complexes may differ, whereby CoV1 RBD may generate strong signaling that favored T_H_1 differentiation to the detriment of T_FH_ differentiation (*36*); thereby impairing downstream GC responses and skewing ASC differentiation towards non-LLPC, leading to antibody waning. Recent studies have instead reported slower decline of spike and RBD antibodies in SARS-CoV-1 versus SARS-CoV-2 convalescent patients (*29*). However, the durability of cognate antibody responses upon vaccination versus infection are expected to be different. Consistently, SARS-CoV-2 convalescents displayed significantly more sustained spike– and RBD-specific antibody responses than mRNA vaccinees (*47*, *48*). A possible explanation may be the broader T cell clonal diversity generated from infection, enabling spike– and RBD-specific B cells to receive additional co-stimulatory signals from T_FH_ cells specific to other viral antigens, thus promoting LLPC differentiation and survival, which would translate into more sustained antibody responses (*49*, *50*).

Finally, this work is the first to report on the suitability of the nasal route for the delivery of Clec9A antibody constructs. We demonstrated that nasal Clec9A^OMNI^ booster in Comirnaty mRNA-vaccinated mice produced robust RBD-specific humoral and cellular responses at both the lower and upper respiratory mucosa, and without compromising the corresponding systemic immune responses. Specifically, potent cross-clade nAb and IgA responses, and functional T_RM_ cells were measured in the lung and nasal compartments, which represent strong correlates of protection against infection and transmission (*51–53*). This was in sharp contrast to the limited induction of respiratory immunity seen upon Comirnaty bivalent mRNA intramuscular booster, where IgG only were detected in the BALF and NLF, likely the result of serum IgG spillover (*7*). Inducing strong nAb response at mucosal sites is important to prevent viruses from establishing infection. IgA in particular serve as the first line of defense in blocking virus attachment and entry via transcytosis towards the apical surface of the mucosal epithelium (*51–54*). On the other hand, mucosal IgG which are largely localized at the basolateral surface, play a protective role in restricting virus replication and dissemination to adjacent tissues and the systemic circulation via direct neutralization or Fc-receptor mediated effector mechanisms (*54–56*). Mucosal T_RM_ cells orchestrate local immune responses to facilitate efficient virus elimination by providing constant immune surveillance of the airway mucosa and initiating rapid recall responses upon cognate antigen encounter (*51*, *57*, *58*). Importantly, we found that nasal Clec9A^OMNI^ boost induced a subset of T_RM_ cells that were CD49a^+^, which has been associated with enhanced Type 1 cytokine and cytolytic responses that have been shown to be critical for antiviral immunity (*58–60*). Therefore, effector responses from reactivated T_RM_ cells could help accelerate local viral clearance and offer early protection against infection and pulmonary disease (*57*, *58*, *61*).

In conclusion, we demonstrated that boosting Comirnaty mRNA-vaccinated mice with Clec9A^OMNI^ elicited sustained, robust, cross-clade, protective immune responses. Given that nAb represents a strong correlate of protection against infection, the ability to generate broadly nAb and cross-reactive B cells against both clade 1a and 1b sarbecoviruses makes Clec9A^OMNI^ an attractive pan-sarbecovirus vaccine candidate (*62*, *63*). Furthermore, successful induction of robust and durable mucosal immunity upon nasal boosting represents a gamechanger that is not only expected to limit virus infection and transmission but may also curb the evolution of new escape variants (*64*, *65*).

As a plug-and-go platform, Clec9A-based constructs can be rapidly updated or engineered in response to newly emerging threats. Moreover, the availability of large scale mAb manufacturing processes worldwide facilitates cost effective production and rapid deployment of Clec9A-targeting vaccines during pandemics compared to other DC-targeting platforms (*66*, *67*). Importantly, human Clec9A-expressing cDC1 subset was found within numerous tissues and mucosae at constant numbers throughout the human lifespan (*68*). Our previous work in humanized mice and non-human primates (*20*, *69*) supports that the Clec9A-targeting vaccine strategy can be readily translated to human populations.

## MATERIALS AND METHODS

### Study design

The goal of this study was to evaluate the ability of Clec9A-RBD booster in the context of COVID19 mRNA vaccine (Comirnaty, Pfizer-BioNTech)-induced pre-existing SARS-CoV2 immunity to trigger cross-clade, durable and protective systemic and mucosal (respiratory) immune responses. mRNA bivalent Comirnaty (BC) (Pfizer-BioNTech) booster immunization was used as benchmark. Sample sizes were determined based on previous experience in immune response studies and viral challenge to achieve statistical significance. All samples and data collected were included with no outlier exclusion. Mice were randomly assigned to the various experimental groups, although no specific method of randomization was used. Investigators performing the experiments and analyses were not blinded. Unless otherwise stated in the figure legends, all *in vivo* experiments and *ex vivo* assays were repeated twice independently with 4-7 mice per group and technical duplicates per biological sample.

### Cell lines and live SARS-CoV-2 virus

ACE2-overexpressing human lung alveolar A549 (A549-ACE2) cells were maintained in Dulbecco’s modified Eagle’s medium (DMEM) (Thermo Fisher Scientific, 11965-118) containing 10% fetal bovine serum (FBS) (Thermo Fisher Scientific, 10270-106) and 1% penicillin-streptomycin (Pen/Strep) (Thermo Fisher Scientific, 15140122).

Omicron BA.1 SARS-CoV-2 virus (GenBank Accession # OP643601) was isolated in NUSMed BSL-3 Core Facility from clinical samples obtained from the National Centre for Infectious Diseases (NCID) and transferred to NUSMed Pathogen Research BSL-2 Lab for virus amplification and titration as described previously (*15*). Briefly, virus was grown in A549-ACE2 cells and culture supernatant was harvested, clarified and concentrated using Amicon 100 kDa filter unit (Merck Millipore, UFC910096) at two dpi. Plaque assay was then performed where virus-containing suspensions were serially diluted ten-fold and incubated with A549-ACE2 cells for one hour at 37°C 5% CO_2_. The infected monolayers were overlaid with DMEM + 2% FBS medium containing 1.2% microcrystalline cellulose. After three days incubation, cells were stained and fixed with 0.5% crystal violet containing 4% formaldehyde for plaque enumeration. Lung viral titer per mL (PFU/mL) was calculated as follow: [(Plaque count)/(Inoculum volume)] x Dilution factor.

### Clec9A-RBD constructs and mRNA vaccines

To generate Clec9A^XBB^ and Clec9A^CoV1^ constructs, rat IgG2a mAb (10B4) specific against mouse Clec9A was genetically fused to one copy of Omicron XBB.1.5 or SARS-CoV-1 RBD **(Fig. S7)**, via a triple alanine linker at the C-terminal end of each heavy chain. Expression, purity and binding of constructs to murine Clec9A, were validated using previously described approaches (*15*). The Pfizer-BioNTech original Comirnaty mRNA vaccine based on Wuhan-Hu-1 (ancestral) SARS-CoV-2 spike protein, and the bivalent Comirnaty (BC) mRNA vaccine based on ancestral and Omicron BA.4/5 spike proteins, were both approved and obtained from the Ministry of Health Singapore.

### Recombinant proteins and peptides

Purified recombinant SARS-CoV-2 ancestral and XBB.1.5 RBD were produced in-house as described previously (*15*). Purified recombinant SARS-CoV-1 RBD (SPD-S52H6), and biotinylated recombinant SARS-CoV-2 ancestral (SPD-C82E9), XBB.1.5 (SPD-C82Q3), and SARS-CoV-1 RBD (SPD-S82E3), were purchased from ACROBiosystems. RBD peptide pools from SARS-CoV-2; ancestral, XBB.1.5 and JN.1, clade 1b bat and pangolin sarbecoviruses; RaTG13 and Gx-P5L, and clade 1a sarbecoviruses; SARS-CoV-1 and WIV-1, were synthesized by Mimotopes Pte Ltd. Each pool comprised of 53 15-mer peptides overlapping by 11 residues and were reconstituted in DMSO (Sigma-Aldrich, 34869).

### Ethics statement

Five to six-week-old female BALB/c mice were purchased from InVivos and housed under specific pathogen-free conditions in individual ventilated cages. All described *in vivo* experiments were approved by the Institutional Animal Care and Use Committee of NUS under protocol numbers R20-0392 and R24-0190 and performed in accordance with the guidelines of the National Advisory Committee for Laboratory Animal Research. Animal facilities are AAALAAC accredited and licensed by the regulatory body Agri-Food and Veterinary Authority of Singapore. All efforts were made to minimize animal suffering.

### Immunization and viral challenge

Mice were injected twice at three-week interval with 0.05 µg original Comirnaty in a volume of 20 µL (diluted in saline) via the intramuscular (i.m) route. Three months after the second dose, mice were boosted with 0.05 µg BC mRNA vaccine via the i.m route, or with Clec9A-RBD constructs supplemented with 50 µg polyinosinic-polycytidylic acid (poly I:C) (Invivogen, vac-pic), either via the subcutaneous (s.c) or intranasal (i.n) route in a volume of 100 µL and 40 µL respectively (diluted in saline). Unless otherwise specified in figure legends, a dose of 10 µg was used for Clec9A^XBB^ and Clec9A^CoV1^ s.c immunization. For Clec9A^OMNI^, 8 µg Clec9A^XBB^ + 2 µg Clec9A^CoV1^ and 4 µg Clec9A^XBB^ + 1 µg Clec9A^CoV1^ were used for s.c and i.n immunization respectively (unless otherwise specified in figure legends). Non-immunized mice served as a negative control.

For challenge experiments, mice were infected i.n with 10^6^ PFU Omicron BA.1 SARS-CoV-2 in a volume of 40 µL (diluted in saline) at one and six months after booster immunization. Non-boosted mice were also included as a susceptibility control. Mice were euthanized at two dpi, and lung tissues were harvested and placed in 2 mL tubes containing 1.4-mm ceramic beads (Omni International, 19-627D) and DMEM + 2% FBS. Whole lung tissues were homogenized for one minute at 4 m/s using Omni Bead Ruptor 12 (Omni International, 19-050A) and centrifuged for one minute at maximum speed to pellet tissue debris. The supernatant was collected as lung homogenate and subjected to plaque assay for lung viral load measurement as described above. Viral titers were expressed as log PFU/mL lung homogenate.

### Sample and tissue collection and processing

Serum and BALF were collected as described previously (*15*). To obtain NLF, mice were euthanized, and a small incision was made at the trachea to allow insertion of a 22G cannula towards the nose. The nasal cavity was gently flushed with 1 mL PBS and the fluid was collected in an Eppendorf tube placed beneath the mouse nostril. The recovered NLF was centrifuged at 400 g for five minutes at 4°C to pellet cell debris before storing the supernatant at –80°C until further analysis.

Following euthanasia at specified timepoints, spleen, lungs, femur and tibia were harvested and processed into single cell suspensions as described previously (*15*). NALT was harvested by dissecting the lower jaw and tongue to reveal the upper palate. After rinsing with chilled PBS, the upper palate was excised using a no. 11 scalpel blade and gently peeled back with fine forceps. The upper palate was minced into small pieces and digested with 0.1 mg/mL Liberase TL (Merck Millipore, 5401020001) in Roswell Park Memorial Institute (RPMI) 1640 medium (Thermo Fisher Scientific, 22400-105) at 37°C 5% CO_2_ for 45 minutes. Cells were passed through a 70 µm cell strainer, centrifuged and resuspended in complete RPMI (cRPMI; RPMI 1640 + 10% FBS + 1% Pen/Strep) to obtain single cell suspension of NALT.

### Enzyme-Linked Immunosorbent Assay (ELISA)

ELISA plates (Sigma-Aldrich, M9410-1CS) were coated with 3 µg/mL of ancestral, XBB.1.5 or SARS-CoV-1 RBD diluted in PBS and incubated overnight at 4°C. For IgG measurement, mouse sera, BALF and NLF were diluted 1:200, 1:20 and 1:10 respectively, followed by two-fold serial dilutions in 1% BSA/PBS (Sigma-Aldrich, A7906). For IgA measurement, BALF and NLF were used neat. Plates were blocked with 5% BSA/PBS for two hours at room temperature (RT), and washed thrice with PBST (PBS, 0.05% Tween 20) before addition of diluted mouse sera, BALF and NLF. After overnight incubation at 4°C, plates were washed thrice, and horseradish peroxidase (HRP)-conjugated anti-mouse IgG (1:3,000) (Bio-rad, 170-6516) and IgA (1:2,000) (Thermo Fisher Scientific, 62-6720) secondary antibodies diluted in 1% BSA/PBS were added and incubated overnight at 4°C. Subsequently, plates were washed thrice and developed using o-phenylenediamine dihydrochloride (OPD) (P9187; Sigma-Aldrich) and 3,30,5,50-tetramethylbenzidine (TMB) (Thermo Fisher Scientific, 00-4201-56) substrate for IgG and IgA detection respectively. Plates were incubated for ten minutes at RT in the dark and the reaction was quenched with 1 M concentrated sulfuric acid (H_2_SO_4_). Absorbance was measured at 490 nm (OPD) and 450 nm (TMB) respectively using Sunrise absorbance microplate reader. Anti-RBD IgG titers were determined by non-linear regression as the reciprocal of the highest dilution with A_490_ corresponding to three times the A_490_ of blank control wells. Anti-RBD IgA titers were expressed as A_450_ values at 1:2 dilution following normalization against A_450_ of blank control wells.

### Multiplex Surrogate Virus Neutralization Test (sVNT)

Multiplex sVNT was established using the Luminex platform as described previously (*1*, *15*). Briefly, RBDs from 22 sarbecoviruses **(Table S1)** were enzymatically biotinylated and coated on MagPlex-Avidin microspheres. Mouse sera were diluted 1:10 and subjected to three four-fold serial dilutions. BALF was subjected to three two-fold serial dilutions while NLF was undiluted. Equal volume of RBD-coated beads was pre-incubated with diluted mouse sera (final dilution: 1:20, 1:80, 1:320, 1:1,280), BALF (final dilution: 1:2, 1:4, 1:8, 1:16) and NLF (final dilution: 1:2) for 15 minutes at 37°C with agitation. Subsequently, PE-conjugated human ACE2 was added to each well and incubated for an additional 15 minutes at 37°C with agitation. After two washes with 1% BSA/PBS, the final readings were acquired using the MAGPIX system according to the manufacturer’s instructions. Percentage inhibition at each dilution was calculated as follows: [(mean fluorescence intensity (MFI) of 30 negative pre-pandemic samples – individual sample FI)/(MFI of 30 negative pre-pandemic samples)] x 100%. Serum and BALF neutralizing activity were reported as the reciprocal of the highest dilution that resulted in >50% inhibition (NT_50_) while NLF neutralizing activity was reported as percentage inhibition at 1:2 dilution, where a cut-off of 30% inhibition or greater was considered as positive for neutralizing antibodies against the sarbecovirus.

### Enzyme-linked immunosorbent spot (ELISPOT)

ELISPOT was performed to detect antigen-specific IgG and IgA ASC. MultiScreen HTS-IP Filter plates (Merck Millipore, MSIPS4W10) were pre-wetted with 35% ethanol for no longer than 1 minute, washed four times with PBS, and coated with 15 µg/mL purified anti-mouse IgG or IgA (Thermo Fisher Scientific, A16080, 62-6700) diluted in PBS. Following overnight incubation at 4°C, plates were washed four times with PBS and blocked with cRPMI for two hours at RT. The blocking solution was discarded and 10^6^ splenocytes, bone marrow and lung cells were added to each respective well and incubated for 40 hours at 37°C and 5% CO_2_. The cells were subsequently discarded, and plates were washed four times with PBST prior to the addition of 1 µg/mL biotinylated ancestral, XBB.1.5 and SARS-CoV-1 RBD diluted in 1% BSA/PBS. Biotinylated anti-mouse IgG and IgA (Biolegend, 405303, 407004) were included for detection of total IgG and IgA secretion as positive controls. After two hours incubation at RT, plates were washed four times with PBST, and HRP-conjugated streptavidin (SAv) (1:100) (BD Biosciences, 554066) diluted in 1% BSA/PBS was added and incubated for one hour at RT. Plates were washed four times with PBST, twice with PBS, and developed with AEC substrate (BD Biosciences, 551951) for 30 minutes before rinsing ten times with deionized water. Spots were enumerated using Mabtech IRIS ELISPOT and FluoroSPOT reader, and analysis was performed using Mabtech Apex software version 1.1.5.74. Data was expressed as spot forming unit (SFU) per 10^6^ cells.

### FluoroSPOT

Mouse IFN-γ/IL-2/TNF-α FluoroSPOT (Mabtech, X-41A42B45W) was performed to evaluate antigen-specific cellular responses, according to the manufacturer’s protocol. Splenocytes, lung and NALT cells (5×10^5^) were restimulated for 40 hours at 37°C and 5% CO_2_ with 2.5 µg/mL ancestral, XBB.1.5, JN.1, RaTG13, Gx-P5L, SARS-CoV-1 and WIV-1 RBD peptides, supplemented with 0.2 µg/mL kit-provided anti-mouse CD28. DMSO and 50 ng/mL PMA + 1 µg/mL ionomycin cocktail (Biolegend, 423302) were used as negative and positive controls respectively. For analysis of spleen CD4^+^ T cell responses, CD8^+^ T cells were depleted from splenocytes using mouse CD8α (Ly-2) microbeads (Miltenyi Biotec, 130-117-044) prior to performing restimulation. Enumeration and analysis of spots were performed similarly as ELISPOT, and data was expressed as SFU per 5×10^5^ cells.

### Flow cytometry

#### Antigen-specific B cell subsets

RBD-specific switched immunoglobulin (swIg^+^) and GC B cells were detected using biotinylated RBD antigens in combination with fluorophore-conjugated SAv. Biotinylated ancestral, XBB.1.5 and SARS-CoV-1 RBD were individually multimerized at a 4:1 molar ratio with SAv-BV421 and SAv-PE, SAv-BV711, and SAv-APC (BD Biosciences, 563259, 554061, 563262, 554067) respectively. SAv-BV605 and SAv-BB515 (BD Biosciences, 563260, 564453) were used as decoy probes without biotinylated RBD to exclude cells with non-specific binding to SAv. After multimerization, RBD probe master mix for swIg^+^ and GC B cell staining were created by mixing the respective fluorophore-conjugated RBD probes together **(Table S2)** with 5 µM free D-biotin (Thermo Fisher Scientific, B20656) to minimize cross-reactivity between probes, diluted in a 1:1 mixture of Brilliant Stain buffer (BD Biosciences, 563794) and fluorescence-activated cell sorting (FACS) buffer (2% FBS + 1 mM EDTA in PBS). Freshly isolated splenocytes, lung (10^6^ and 5×10^6^ for swIg^+^ and GC B cell analysis respectively) and NALT cells (5×10^5^ for swIg^+^ B cell analysis) were incubated with anti-CD16/CD32 Fc block (BD Biosciences, 553142) and eFluor 780 Fixable Viability Dye (Thermo Fisher Scientific, 65-0865-14) diluted in FACS buffer (1:200 and 1:1,000 respectively) for 20 minutes at 4°C. Cells were washed with FACS buffer and stained with respective decoy probe **(Table S2)** diluted in FACS buffer with 5 µM free D-biotin for 30 minutes at 4°C in the dark. Subsequently, cells were washed and stained with RBD probe master mix for one hour at 4°C in the dark. Thereafter, cells were washed and stained with swIg^+^ or GC B cell surface marker antibodies **(Table S2)** diluted in Brilliant Stain and FACS buffer for 30 minutes at 4°C in the dark.

#### Activation-induced marker (AIM^+^) T_FH_ cells

Splenocytes and lung cells (2×10^6^) were stimulated with 2.5 µg/mL ancestral RBD peptides in the presence of 1 µg/mL purified anti-mouse CD40 and anti-mouse CD154-PE (1:100) (Biolegend, 102802, 106506) at 37°C and 5% CO_2_. DMSO and PMA + Ionomycin cocktail were included as negative and positive controls respectively, similarly as above. After overnight incubation, cells were washed and incubated with anti-CD16/CD32 Fc block and eFluor780 Fixable Viability Dye, similarly as above. Thereafter, cells were washed and stained with AIM^+^ T_FH_ cell surface marker antibodies **(Table S2)**, similarly as above.

#### Tissue-resident memory T (T_RM_) cells

Freshly isolated lung (10^6^) and NALT (5×10^5^) cells were incubated with anti-CD16/CD32 Fc block and eFluor780 Fixable Viability Dye, similarly as above. Cells were washed and stained with T_RM_ cell surface marker antibodies **(Table S2)**, similarly as above.

#### Data acquisition and analysis

Cells were washed and resuspended in FACS buffer after the final staining step prior to data acquisition on the LSRFortessa II X-20 analyzer (BD Biosciences). UltraComp eBeads (Thermo Fisher Scientific, 01-222-42) were used for single colour compensation controls. Gating strategies and representative analysis results for antigen-specific B cell subsets **(Fig. S8, S11, S12)**, AIM^+^ T_FH_ **(Fig. S9, S11, S13)** and T_RM_ cells **(Fig. S9, S13)** can be found in the supplementary materials.

### FACS sorting of long-lived plasma cell (LLPC) and non-LLPC ASC subsets

Freshly isolated bone marrow cells (6-8×10^7^) were incubated with anti-CD16/CD32 Fc block and eFluor780 Fixable Viability Dye, similarly as above. Cells were washed and stained with ASC subset surface marker antibodies **(Table S2)** diluted in Brilliant Stain and FACS buffer for 30 minutes at 4°C. Thereafter, cells were washed, resuspended in FACS buffer, and sorted for LLPC and non-LLPC subsets using the FACSAria Fusion™ Cell Sorter (BD Biosciences) and gating strategy indicated in Fig. S10. Total and anti-RBD IgG secretion from each ASC subset were assessed by ELISPOT as described above, which quantified frequency of total and anti-RBD IgG-secreting cells. The number of input ASC were 1,000-2,000 and 10,000-20,000 for total and anti-RBD IgG detection respectively. Data was normalized and expressed as % anti-RBD IgG ASC out of total IgG ASC for each subset, which was calculated as follow: (SFU_RBD IgG_)/(SFU_Total IgG_) x 100%.

### Statistical analysis

All data were plotted and analyzed using PRISM GraphPad Version 10, and statistical significance was assigned when p-value < 0.05 (*), <0.01 (**) and <0.001 (***). Details about the number of animals, descriptive statistics reported, statistical tests and comparisons used, are outlined in the figure legends. In brief, non-parametric two-tailed Mann Whitney test (two groups) and Kruskal-Walis test (more than two groups) with Dunnett’s correction for multiple comparisons were used for unpaired group analysis, while Wilcoxon matched-pairs signed rank test (two groups) and Friedman test with Dunnett’s correction for multiple comparisons (more than two groups) were used for paired group analysis.

## List of Supplementary Materials

-Fig S1 to S13

-Tables S1 to S2

## Acknowledgements

We thank A/P Paul Hutchinson and his team from the flow cytometry core facility at Life Sciences Institute, NUS for their assistance.

## Author contributions

-Performed the experiments: NYZC; WCY, PST, XQ, KMT, KP, WYT, SYYM –Provided inputs: PAM; CWT; MHL; SA. –Wrote the manuscript: NYZC; SA.

## Data and materials availability

The authors will make all reagents and materials available to the scientific community upon request. Clec9A constructs are available under MTA with Monash University.

## SUPPLEMENTARY MATERIALS

**Figure S1.**
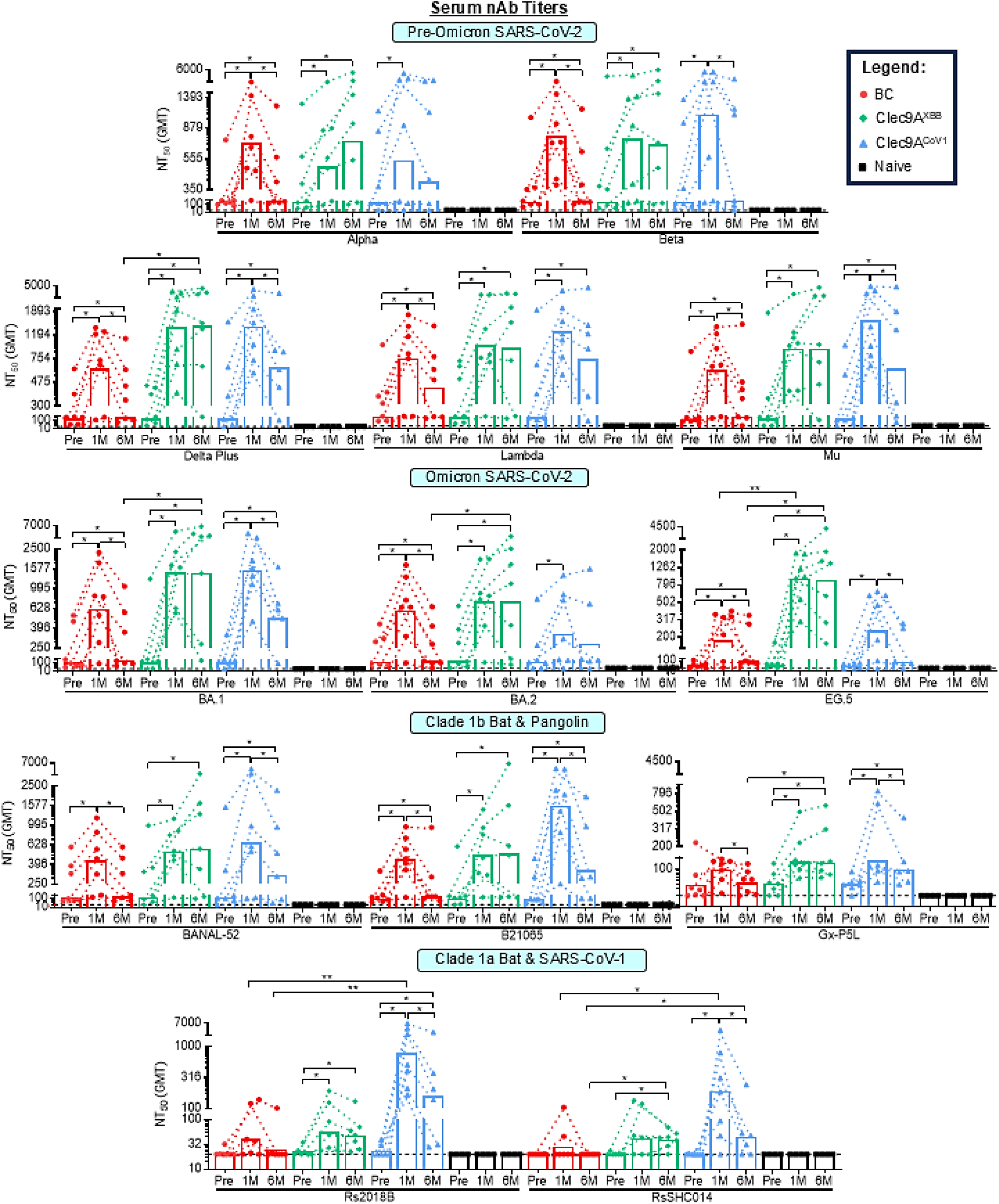
Serum neutralizing antibody titers upon systemic booster immunization with Clec9A^XBB^, Clec9A^CoV1^ or BC mRNA vaccine. Five to six-week-old BALB/c mice (n = 6-7 per group) were immunized twice three weeks apart (0.05 µg per dose; i.m) with Pfizer-BioNTech original Comirnaty mRNA vaccine. Three months after the last immunization dose, mice were boosted either with Pfizer-BioNTech BA.4/5 bivalent Comirnaty (BC) mRNA vaccine (0.05 µg; i.m), Clec9A^XBB^ (10 µg adjuvanted with 50 µg poly I:C; s.c) or Clec9A^CoV1^ (10 µg adjuvanted with 50 µg poly I:C; s.c). A control group of non-immunized mice (naïve) was also included for baseline. The serum nAb titers (NT_50_) against 13 sarbecoviruses representative of clades 1b and 1a were determined by multiplex sVNT at one week pre-boost (Pre), one– and six months post-boost (1M, 6M). Symbols represent individual animals and data shown are geometric means. Statistical analysis: Non-parametric two-tailed Kruskal Walis test with Dunnett’s correction for multiple comparisons and Friedman test with Dunnett’s correction for multiple comparisons. *p < 0.05, **p < 0.01.

**Figure S2.**
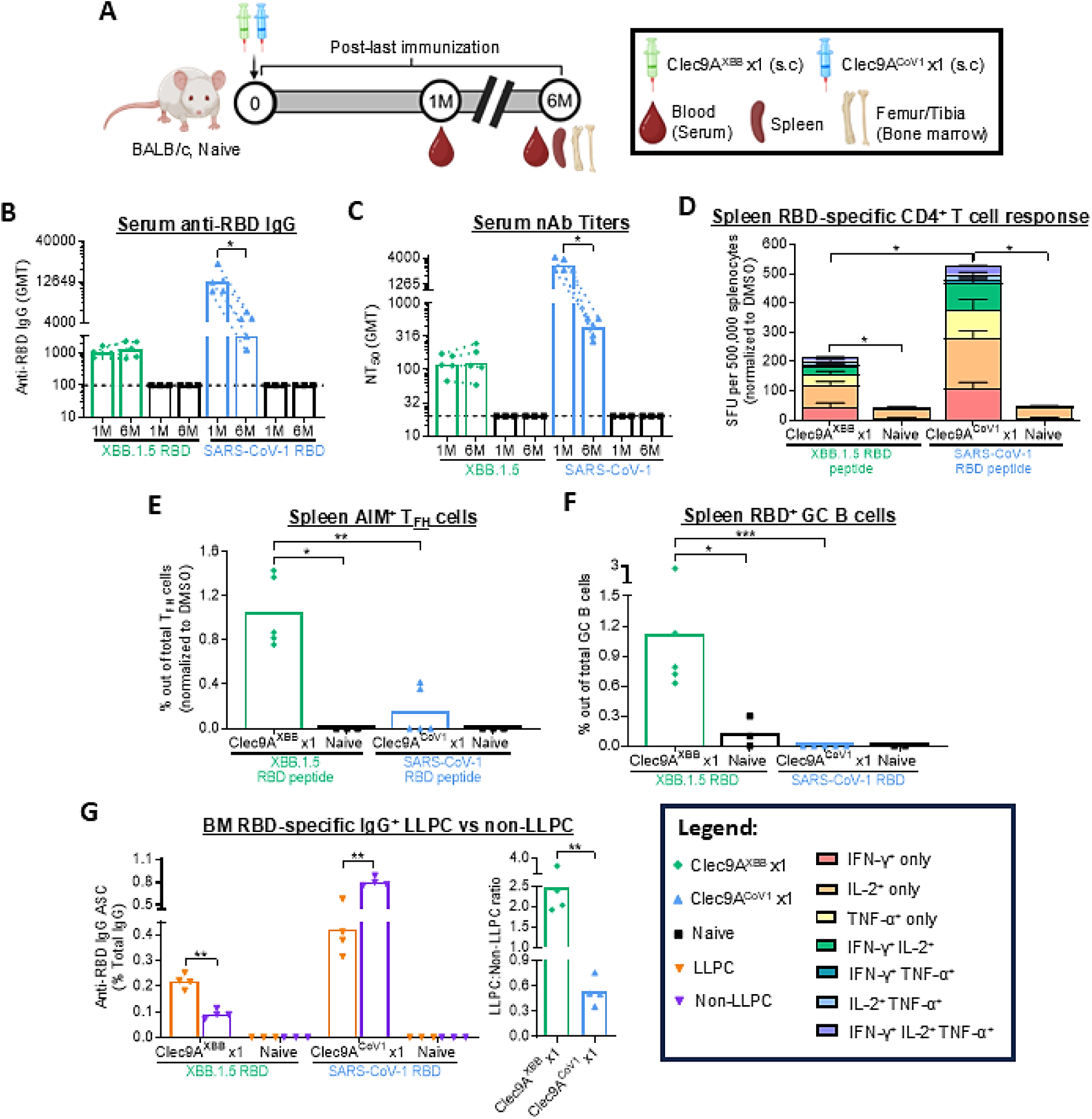
Durability of RBD-specific immune responses upon single shot immunization with Clec9A^XBB^ or Clec9A^CoV1^ in naïve mice. **A)** Immunogenicity experiment design. Five to six-week-old BALB/c mice (n = 4-5 per group) were immunized with a single dose (2 µg adjuvanted with 50 µg poly I:C; s.c) of Clec9A^XBB^ or Clec9A^CoV1^. **(B)** Blood was collected one– and six months after immunization. Serum IgG titers against XBB.1.5 and SARS-CoV-1 RBD were determined by ELISA. The dashed line represents the LOD at 100. **(C)** Serum nAb titers (NT_50_) against XBB.1.5 and SARS-CoV-1 at one– and six months post-immunization were determined by sVNT. **(D)** Frequency of IFN-γ, IL-2 and/or TNF-α secreting CD4^+^-enriched splenocytes at six months post-immunization was determined by FluoroSPOT upon re-stimulation with XBB.1.5 and SARS-CoV-1 RBD peptides. SFU = spot forming units. **(E, F)** Percentage of **(E)** AIM^+^ T_FH_ and **(F)** RBD^+^ GC B cells in spleen at six months post-immunization was determined by flow cytometry. **(G)** Frequency of BM RBD-specific IgG^+^ LLPC and non-LLPC (normalized to total IgG) at six months post-immunization was determined by B cell ELISPOT. **(B, C, E-G)** Symbols represent individual animals and data shown are **(B, C)** geometric means and **(D-G)** means ± **(D)** SD. Statistical analysis: Non-parametric two-tailed **(B, C)** Wilcoxon matched-pairs signed rank test and **(D-G)** Mann-Whitney test. *p < 0.05, **p < 0.01, ***p < 0.001.

**Figure S3.**
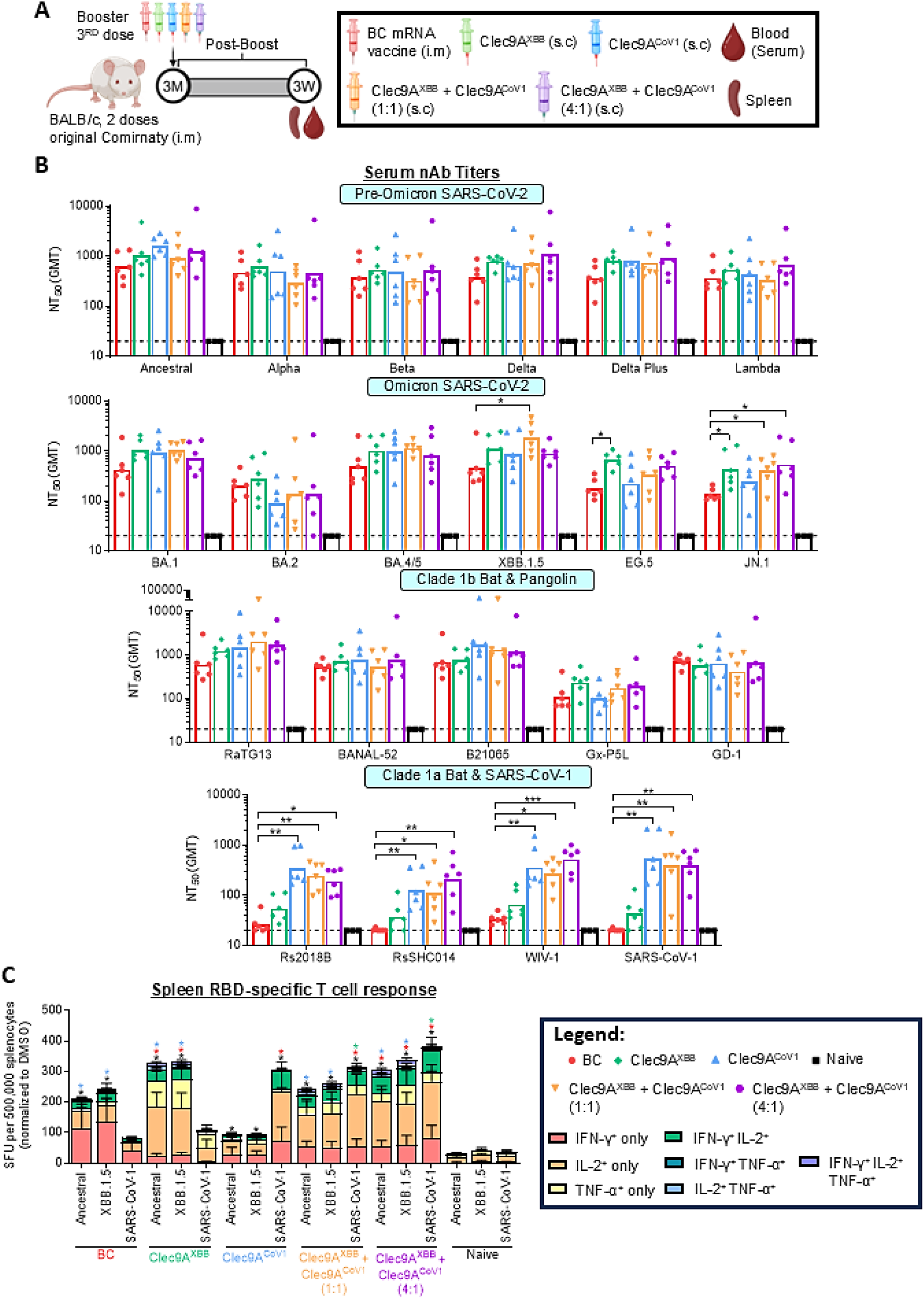
Dose optimization of [Clec9A^XBB^ + Clec9A^CoV1^] booster immunization. **(A)** Immunogenicity experiment design. Five to six-week-old BALB/c mice (n = 5-6 per group) were immunized twice three weeks apart (0.05 µg per dose; i.m) with original Comirnaty mRNA vaccine. Three months after the second immunizing dose, mice were boosted with either BC mRNA vaccine (0.05 µg; i.m), Clec9A^XBB^ (10 µg), Clec9A^CoV1^ (10 µg), [Clec9A^XBB^ + Clec9A^CoV1^] (5 µg + 5 µg) or [Clec9A^XBB^ + Clec9A^CoV1^] (8 µg + 2 µg). All Clec9A-based formulations were adjuvanted with 50 µg poly I:C and administered s.c. At three weeks post-boost, blood was collected, and mice were euthanized for spleen harvesting. **(B)** Serum nAb titers (NT_50_) against 21 sarbecoviruses from clades 1a and 1b were determined by multiplex sVNT. **(C)** Frequency of IFN-γ, IL-2 and/or TNF-α secreting splenocytes was determined by FluoroSPOT upon re-stimulation with ancestral SARS-CoV-2, XBB.1.5 and SARS-CoV-1 RBD peptides. SFU = spot forming units. Data shown are **(B)** geometric means and **(C)** means ± SD. Statistical analysis: Non-parametric two-tailed Kruskal Walis test with Dunnett’s correction for multiple comparisons. *p < 0.05, **p < 0.01, ***p < 0.001.

**Figure S4.**
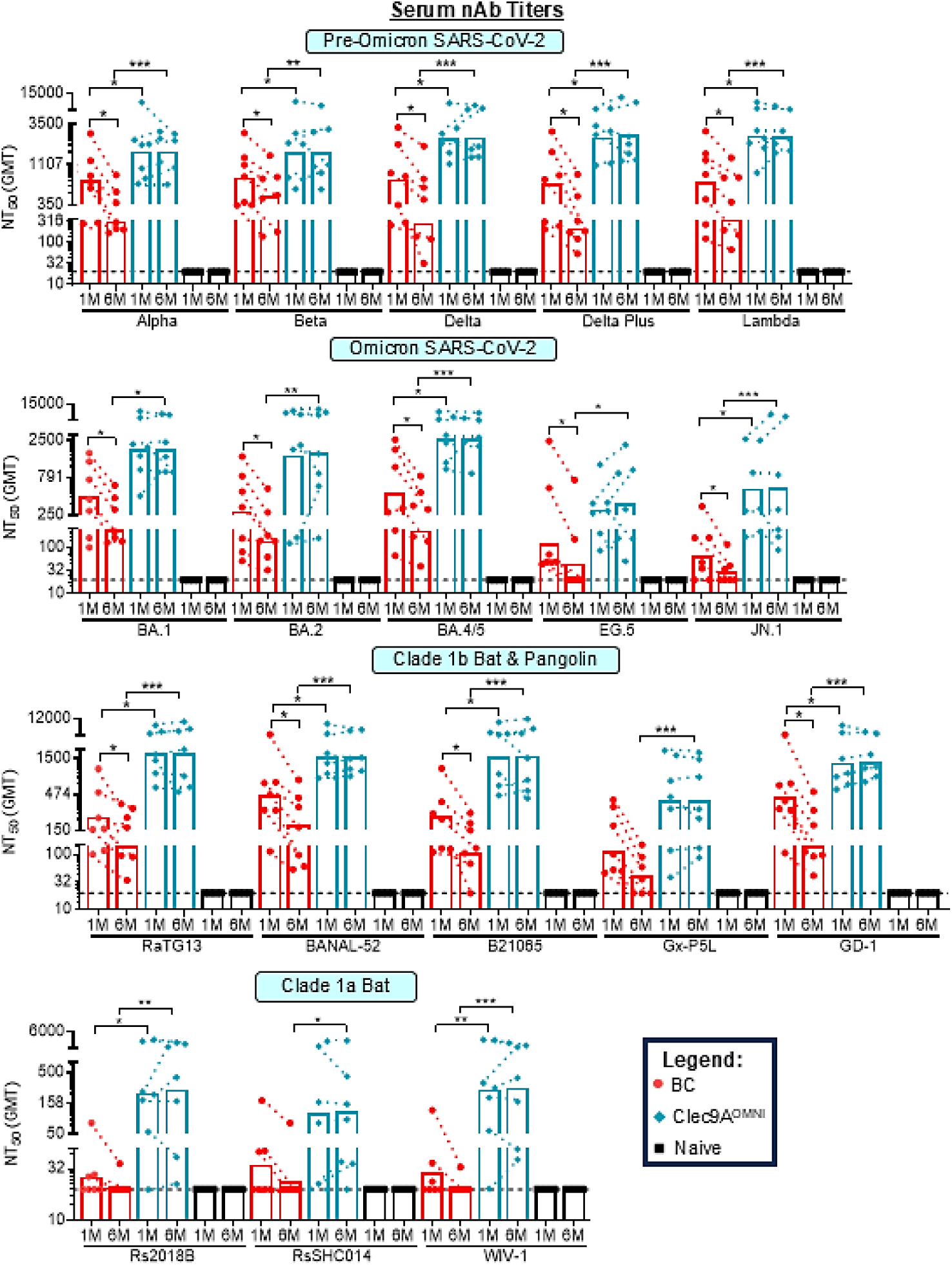
Serum neutralizing antibody responses upon systemic booster immunization with Clec9A^OMNI^ versus BC mRNA vaccine. Five to six-week-old BALB/c mice (n = 5-6 per group) were immunized twice three weeks apart (0.05 µg per dose; i.m) with original Comirnaty mRNA vaccine. Three months after the second immunization dose, mice were boosted with either BC mRNA vaccine (0.05 µg; i.m), or Clec9A^OMNI^ (8 µg Clec9A^XBB^ + 2 µg Clec9A^CoV1^ adjuvanted with 50 µg poly I:C; s.c). Serum nAb titers (NT_50_) against 18 sarbecoviruses from clades 1a and 1b were determined at one– and six months post-boost by multiplex sVNT. Symbols represent individual animals and data shown are geometric means. Statistical analysis: Non-parametric two-tailed Mann-Whitney test, and Friedman test with Dunnett’s correction for multiple comparisons. *p < 0.05, **p < 0.01, ***p < 0.001.

**Figure S5.**
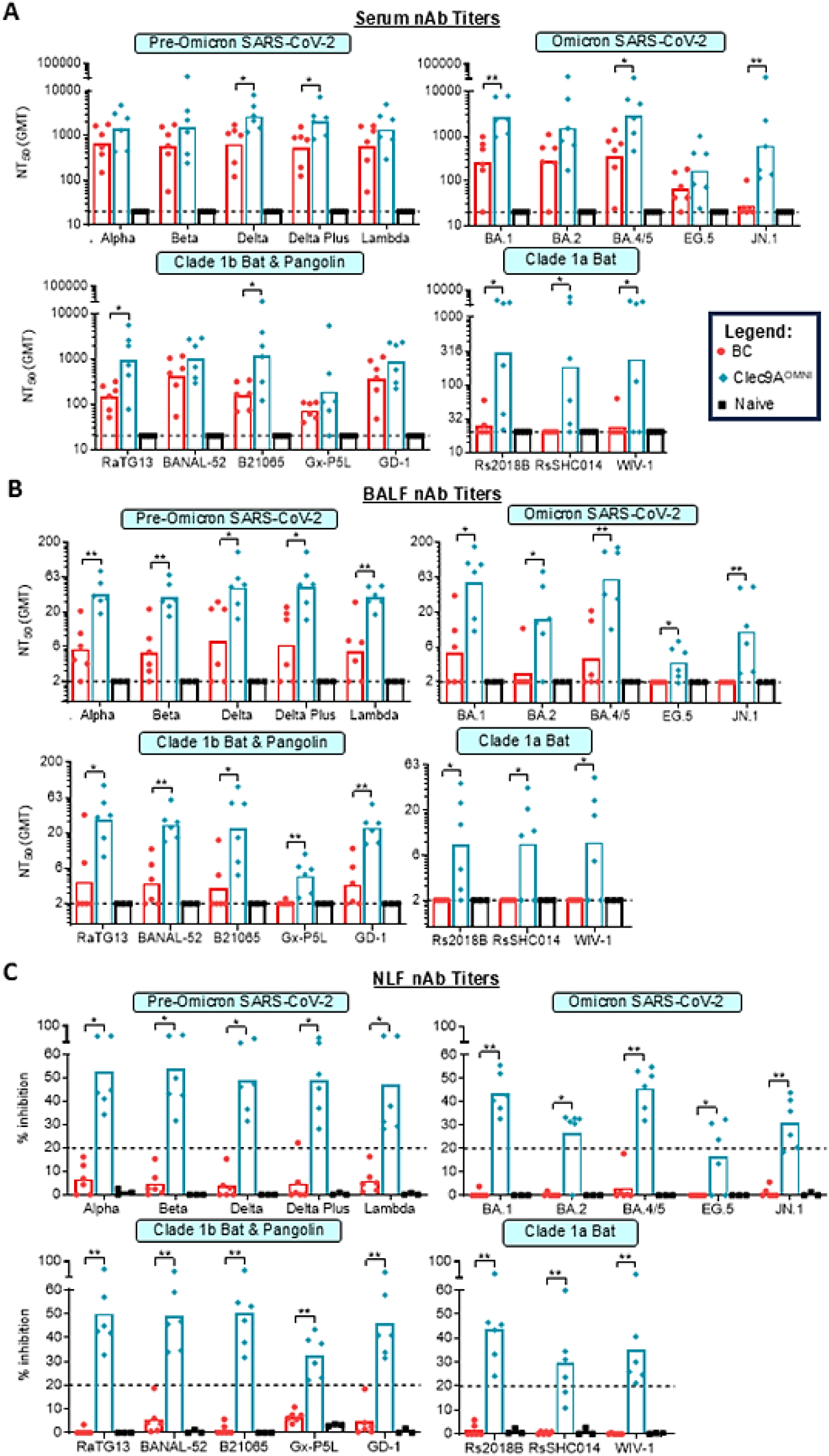
Neutralizing antibody responses upon nasal booster immunization with Clec9A^OMNI^ versus systemic booster immunization with BC mRNA vaccine. Five to six-week-old BALB/c mice (n = 5-6 per group) were immunized twice three weeks apart with original Comirnaty mRNA vaccine (0.05 µg per dose; i.m). Three months after the second immunization dose, mice were boosted with either BC mRNA vaccine (0.05 µg; i.m), or Clec9A^OMNI^ (4 µg Clec9A^XBB^ + 1 µg Clec9A^CoV1^ adjuvanted with 50 µg poly I:C; i.n). **(A)** Serum, **(B)** BALF and **(C)** NLF nAb titers (serum, BALF: NT_50_, NLF: % inhibition at 1:2 dilution) against 18 sarbecoviruses from clades 1a and 1b were determined at one-month post-boost by multiplex sVNT. **(A-C)** Symbols represent individual animals and data shown are **(A, B)** geometric means and **(C)** means. Statistical analysis: Non-parametric two-tailed Mann-Whitney test. *p < 0.05, **p < 0.01.

**Figure S6.**
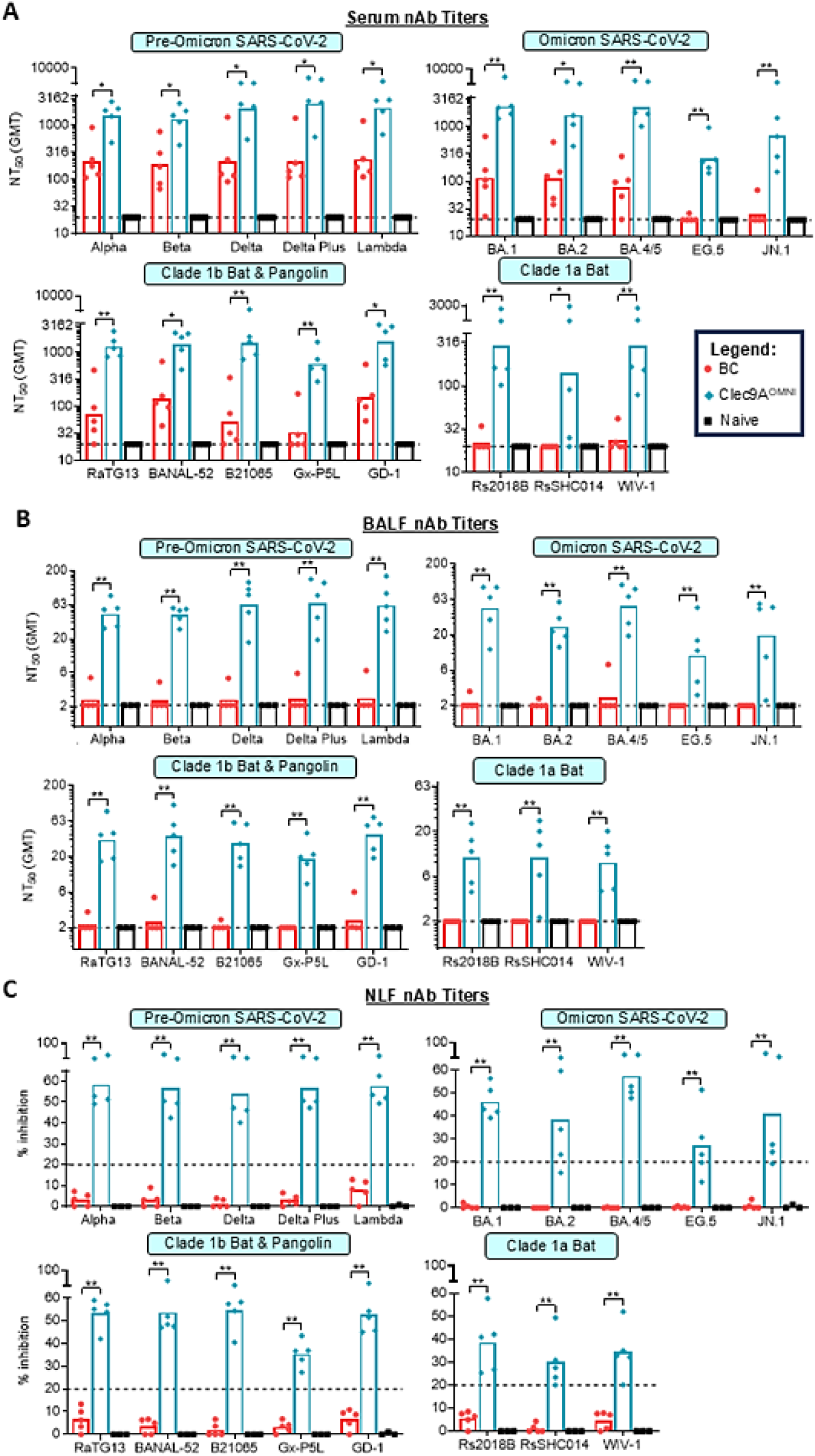
Long-term neutralizing antibody responses upon nasal booster immunization with Clec9A^OMNI^ versus systemic booster immunization with BC mRNA vaccine. Five to six-week-old BALB/c mice (n = 5-6 per group) were immunized twice three weeks apart with original Comirnaty mRNA vaccine (0.05 µg per dose; i.m). Three months after the second immunization dose, mice were boosted with either BC mRNA vaccine (0.05 µg; i.m), or Clec9A^OMNI^ (4 µg Clec9A^XBB^ + 1 µg Clec9A^CoV1^ adjuvanted with 50 µg poly I:C; i.n). **(A)** Serum, **(B)** BALF and **(C)** NLF nAb titers (serum, BALF: NT_50_, NLF: % inhibition at 1:2 dilution) against 18 sarbecoviruses from clades 1a and 1b were determined at six months post-boost by multiplex sVNT. **(A-C)** Symbols represent individual animals and data shown are **(A, B)** geometric means and **(C)** means. Statistical analysis: Non-parametric two-tailed Mann-Whitney test. *p < 0.05, **p < 0.01.

**Figure S7.**
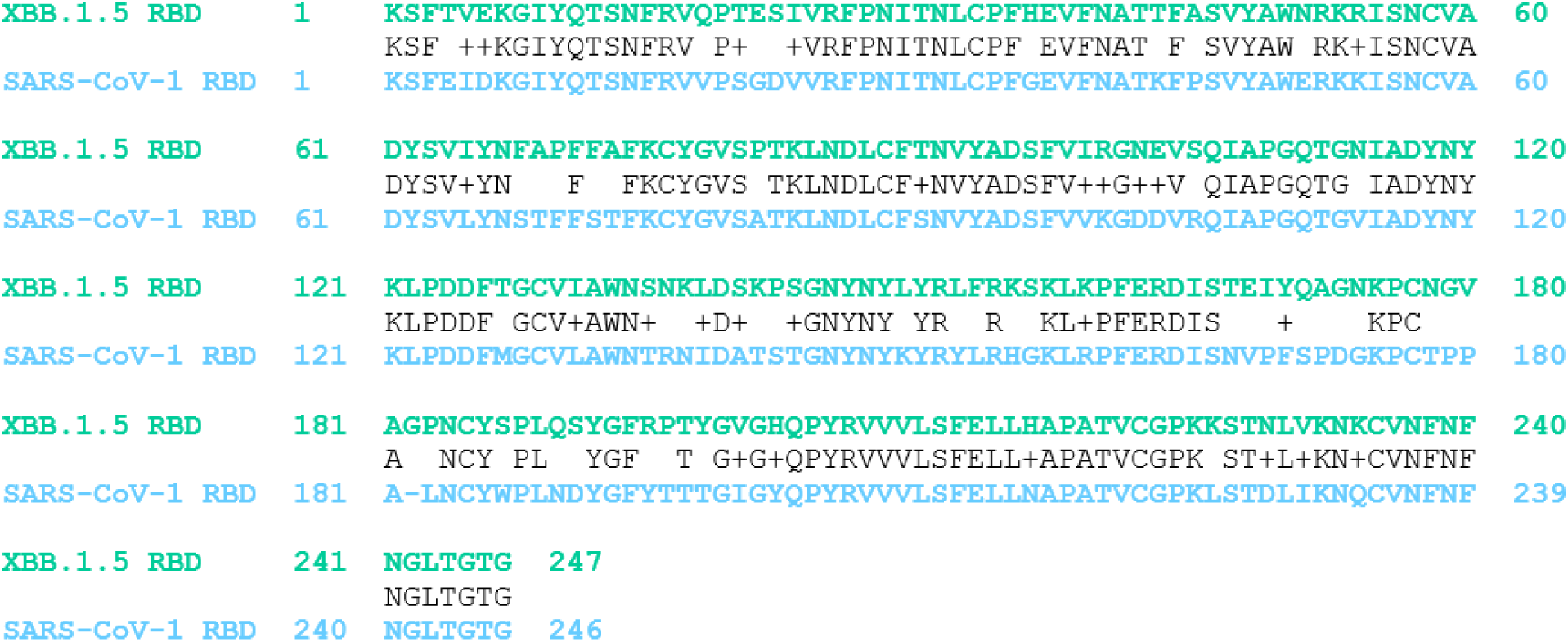
BLAST amino acid sequence alignment for RBD antigens (XBB.1.5 and SARS-CoV-1) used for expression of Clec9A-RBD constructs.

**Figure S8.**
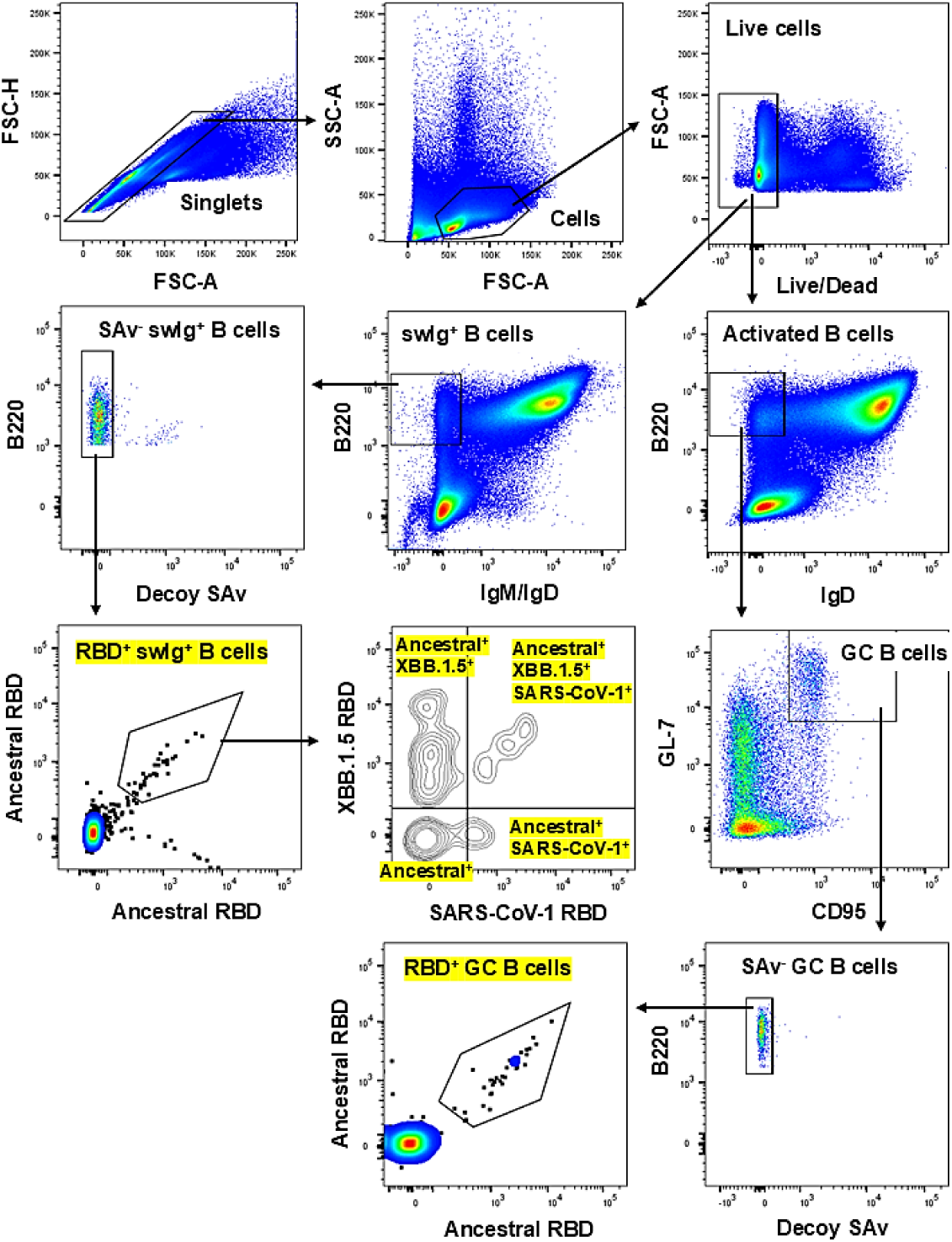
Gating strategy to identify antigen-specific swIg^+^ and GC B cell subsets. Single cells were first identified via FSC-H/FSC-A and SSC-A/FSC-A. Dead cells were excluded with eFluor780 Fixable Viability Dye, and swIg^+^ (B220^+^ IgD^-^ IgM^-^) and activated (B220^+^ IgD^-^) B cells were identified from live cell population (Live/Dead^-^). For the former, swIg^+^ B cells that bind non-specifically to SAv were excluded using decoy SAv probe, and RBD-specific swIg^+^ B cells were identified from the SAv^-^ swIg^+^ B cell population via a double discrimination gate where cells must be double positive for both ancestral RBD-BV421 and –PE to be considered as antigen-specific. Cross-reactivity to XBB.1.5 and SARS-CoV-1 RBD were further determined from the gated total RBD^+^ swIg^+^ B cell population. For the latter, GC B cells (GL-7^+^ CD95^+^) were identified from the activated B cell population, and those binding non-specifically to SAv were excluded using decoy SAv probe. RBD-specific GC B cells were subsequently identified from the SAV^-^ GC B cell population via double discrimination gating.

**Figure S9.**
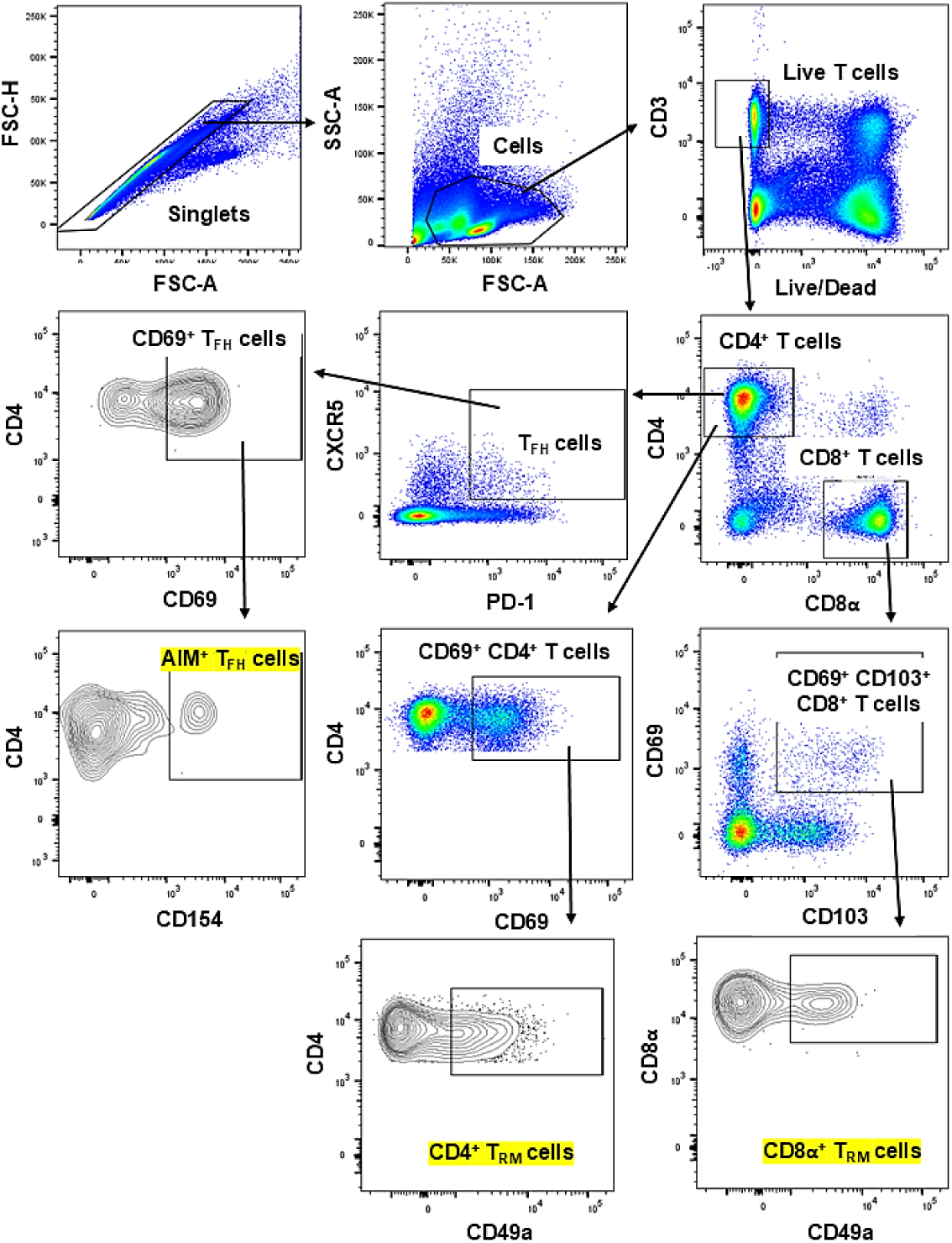
Gating strategy to identify AIM^+^ T_FH_ and T_RM_ cells. Single cells were first identified via FSC-H/FSC-A and SSC-A/FSC-A. Dead cells were excluded with eFluor780 Fixable Viability Dye, and CD4^+^ and CD8^+^ T cells were identified from the total live T cell (Live/Dead^-^ CD3^+^) population. AIM^+^ T_FH_ cells were examined by first identifying total T_FH_ cells (CXCR5^+^ PD-1^+^) from the CD4^+^ T cell population, followed by gating on those that are double positive for both CD69 and CD154 AIM. T_RM_ cells in lung and NALT tissues were identified from total CD4^+^ and CD8^+^ T cells by gating into CD69^+^ and CD69^+^ CD103^+^ populations respectively, followed by further gating into cells that were also CD49a^+^ (CD4^+^: CD69^+^ CD49a^+^, CD8^+^: CD69^+^ CD103^+^ CD49a^+^).

**Figure S10.**
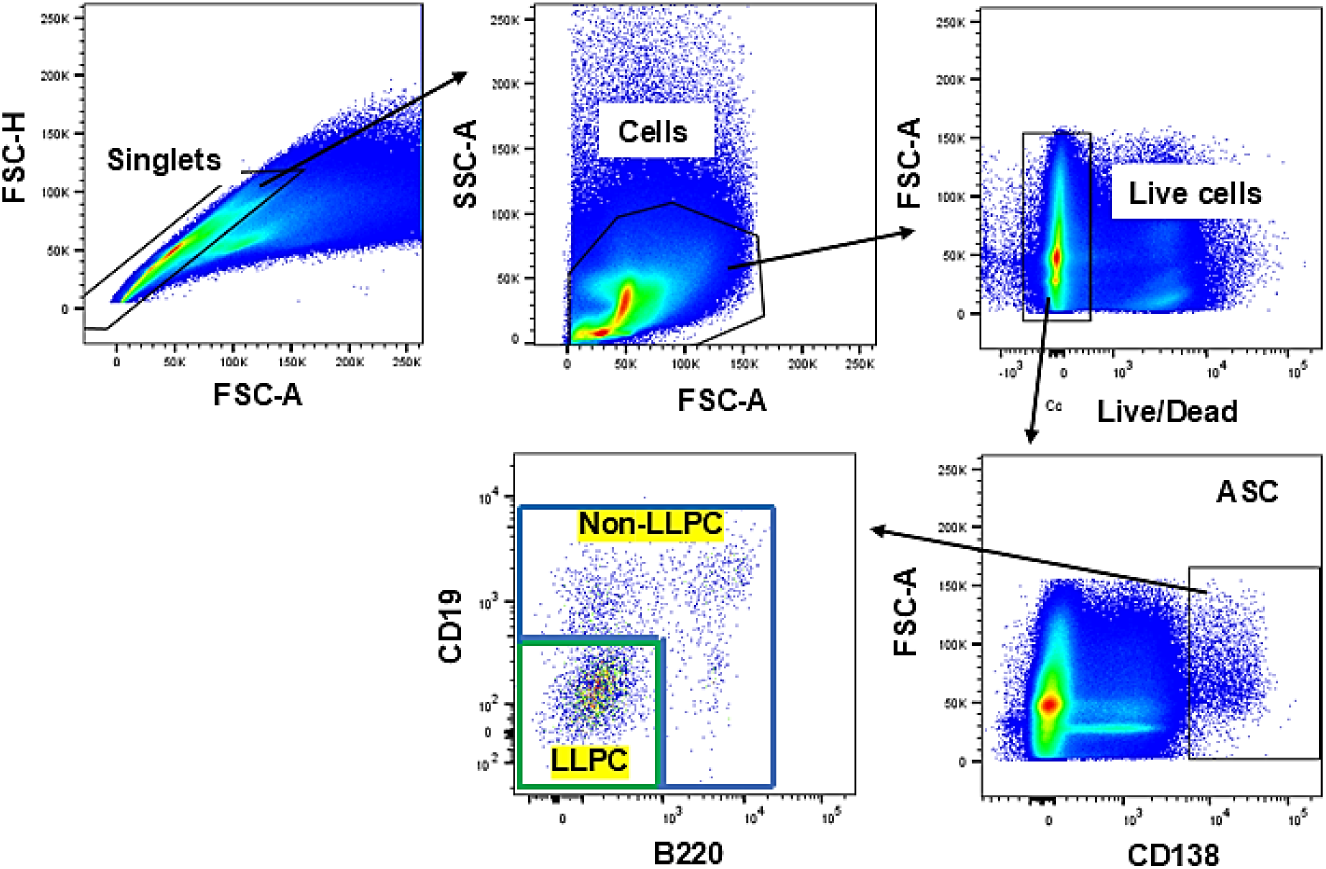
Gating strategy for sorting bone marrow LLPC and non-LLPC ASC subsets. Single cells were first identified via FSC-H/FSC-A and SSC-A/FSC-A. Dead cells were excluded with eFluor780 Fixable Viability Dye, and ASC population was identified as live cells that highly express CD138 (Live/Dead^-^ CD138^hi^). Boolean gating was subsequently applied to identify LLPC and non-LLPC ASC subsets based on their expression of B220 and CD19; LLPC = CD138^hi^ B220^lo^ CD19l° (green), non-LLPC = inverse of LLPC gating (blue).

**Figure S11.**
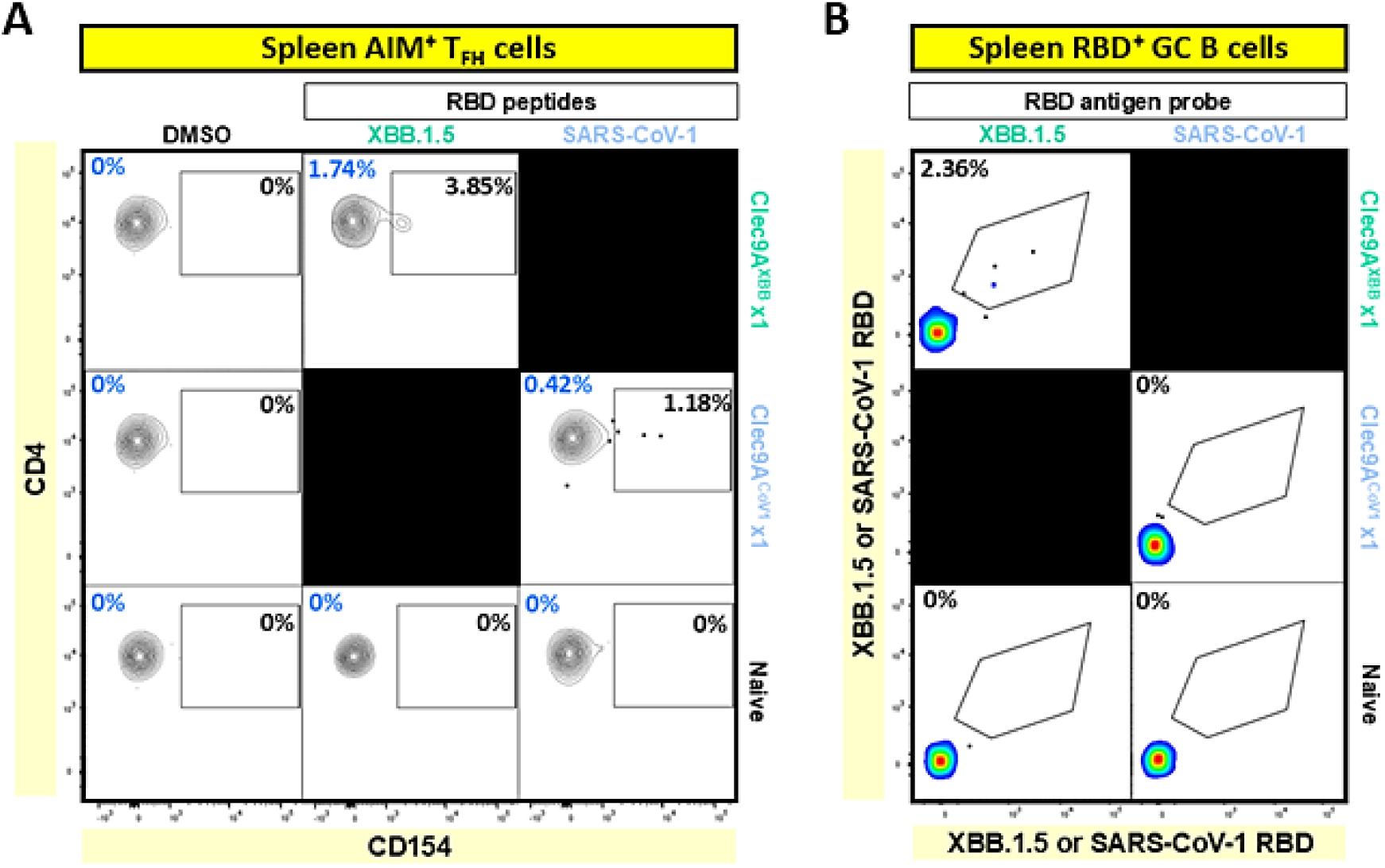
Representative flow cytometric analysis results of AIM^+^ T_FH_ and RBD^+^ GC B cells following single dose Clec9A^XBB^ or Clec9A^CoV1^ immunization. **(A)** Representative plots of spleen AIM^+^ T_FH_ cells at six months post-boost. Values indicated in black and blue represent percentage AIM^+^ T_FH_ cells out of CD69^+^ T_FH_ cells and total T_FH_ cells respectively. **(B)** Representative plots of RBD-specific spleen GC B cells at six months post-boost.

**Figure S12.**
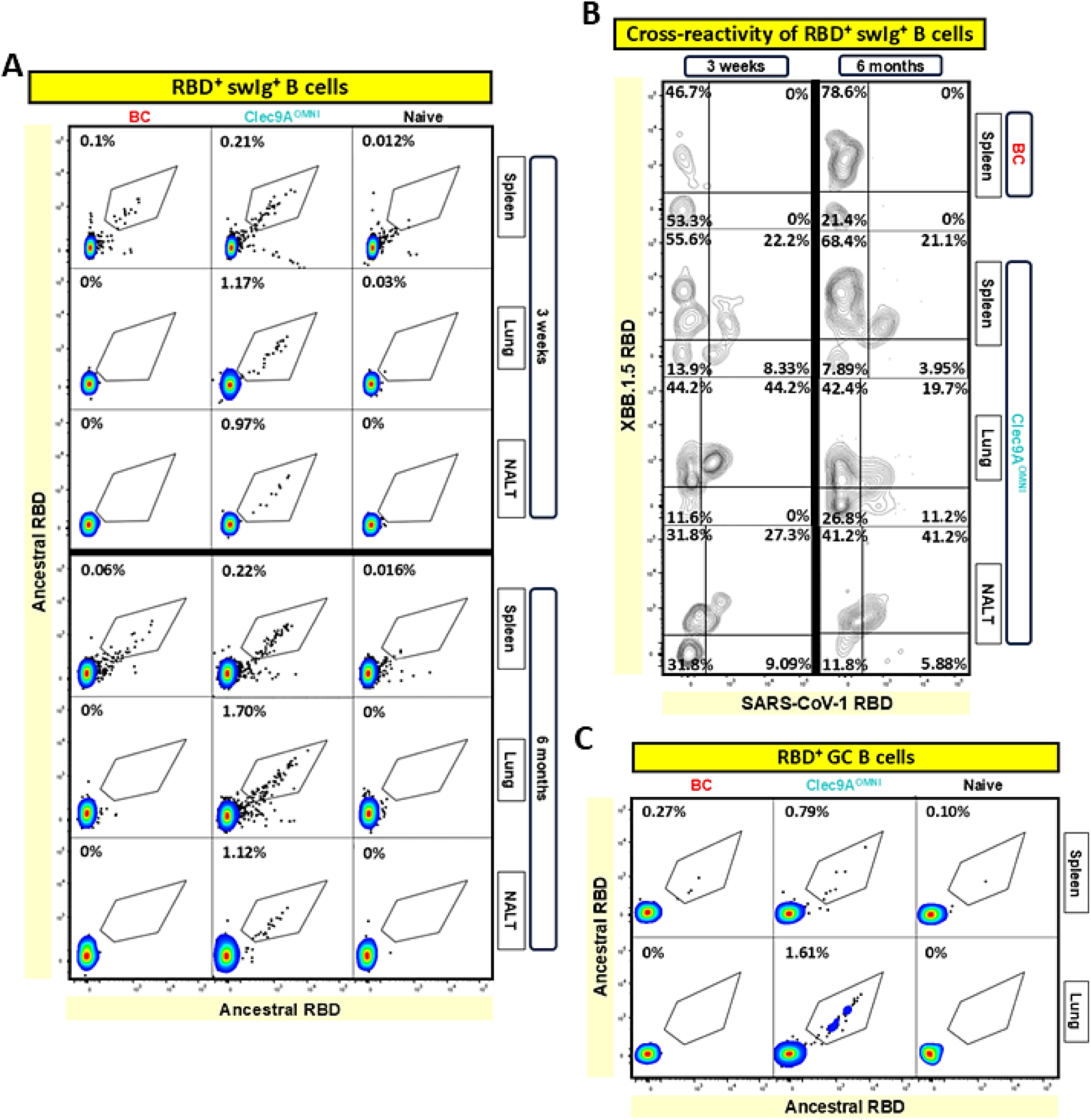
Representative flow cytometric analysis results of antigen-specific B cell subsets following BC mRNA vaccine and Clec9A^OMNI^ booster immunization. **(A, B)** Representative plots of ancestral SARS-CoV-2 RBD-specific spleen, lung and NALT **(A)** swIg^+^ B cells and their **(B)** cross-reactivity to XBB.1.5 and SARS-CoV-1 RBD, at three weeks and six months post-boost. **(C)** Representative plots of RBD-specific spleen and lung GC B cells at six months post-boost.

**Figure S13.**
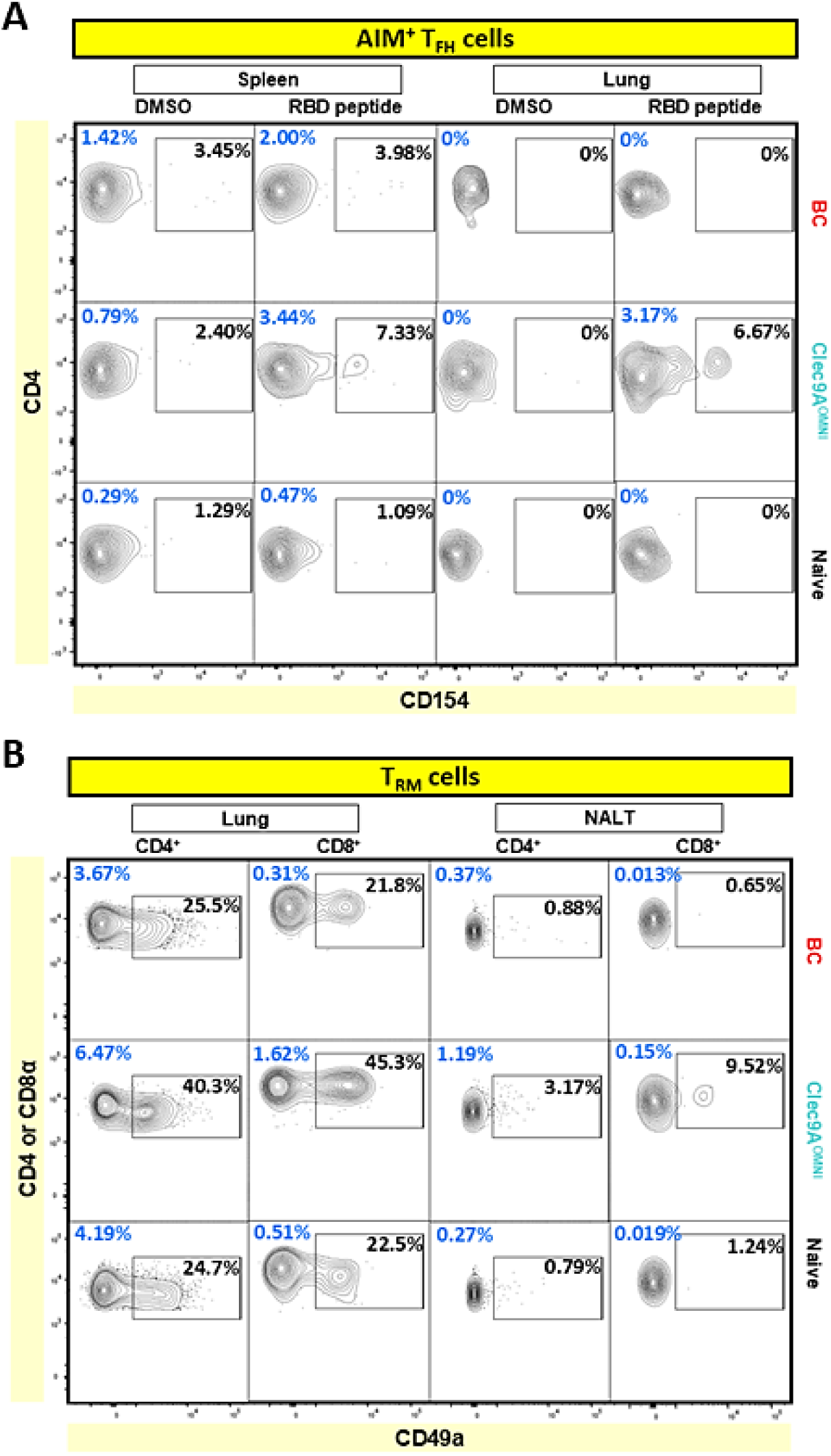
Representative flow cytometric analysis results of AIM^+^ T_FH_ and T_RM_ cells following BC mRNA vaccine and Clec9A^OMNI^ booster immunization. **(A)** Representative plots of spleen and lung AIM^+^ T_FH_ cells at six months post-boost. Values indicated in black and blue represent percentage AIM^+^ T_FH_ cells out of CD69^+^ T_FH_ cells and total T_FH_ cells respectively. **(B)** Representative plots of lung and NALT CD4^+^ and CD8^+^ T_RM_ cells at one-month post-boost. Values indicated in black and blue represent percentage T_RM_ cells out of CD69^+^ CD4^+^ or CD69^+^ CD103^+^ CD8^+^ T cells, and total CD4^+^ or CD8^+^ T cells respectively.

**Table S1.**
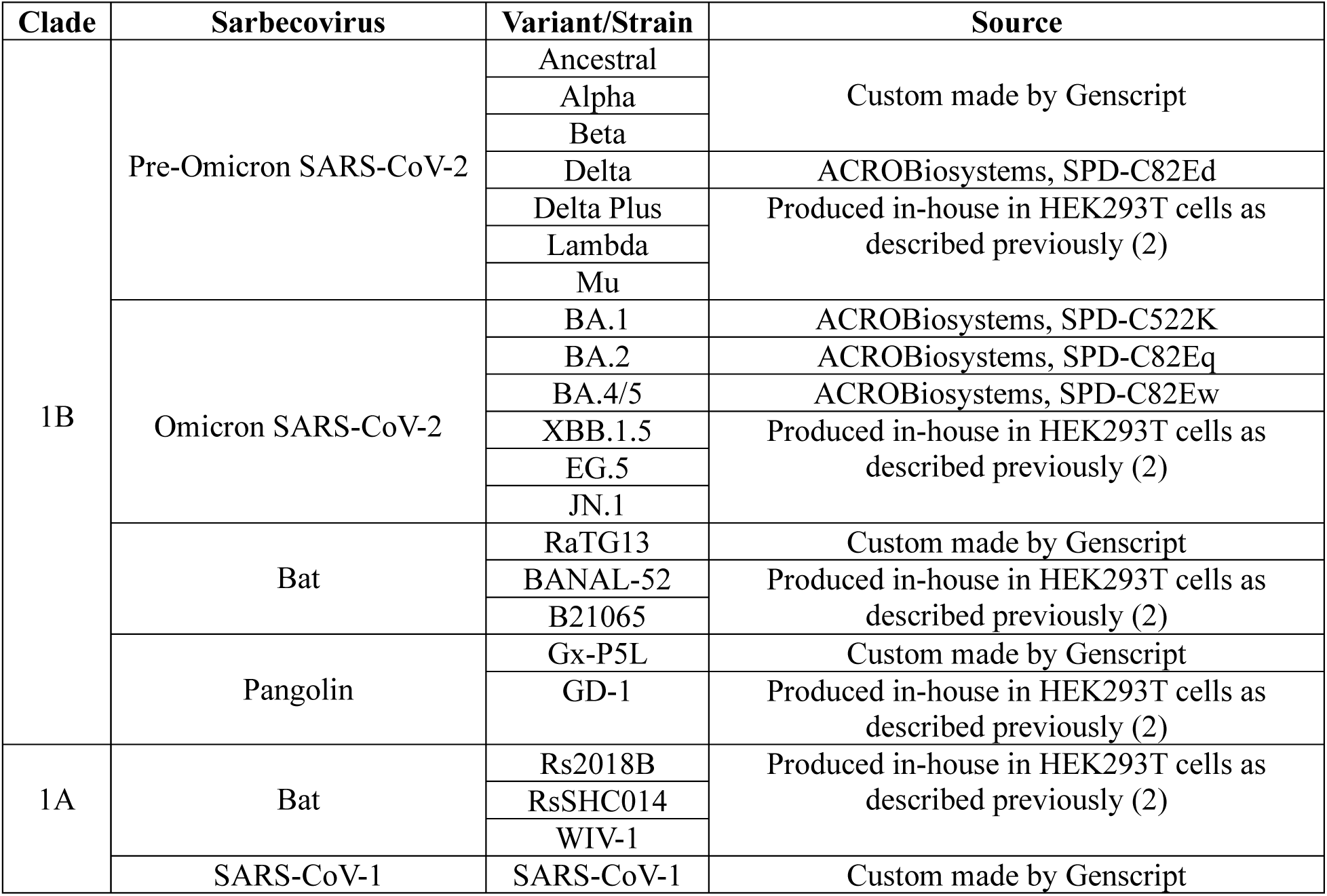
Sarbecovirus RBD antigens used in the Multiplex sVNT.

| Clade | Sarbecovirus | Variant/Strain | Source |
| --- | --- | --- | --- |
| 1B | Pre-Omicron SARS-CoV-2 | Ancestral | Custom made by Genscript |
|  |  | Alpha |  |
|  |  | Beta |  |
|  |  | Delta | ACROBiosystems, SPD-C82Ed |
|  |  | Delta Plus | Produced in-house in HEK293T cells as described previously (2) |
|  |  | Lambda |  |
|  |  | Mu |  |
|  | Omicron SARS-CoV-2 | BA.1 | ACROBiosystems, SPD-C522K |
|  |  | BA.2 | ACROBiosystems, SPD-C82Eq |
|  |  | BA.4/5 | ACROBiosystems, SPD-C82Ew |
|  |  | XBB.1.5 | Produced in-house in HEK293T cells as described previously (2) |
|  |  | EG.5 |  |
|  |  | JN.1 |  |
|  | Bat | RaTG13 | Custom made by Genscript |
|  |  | BANAL-52 | Produced in-house in HEK293T cells as described previously (2) |
|  |  | B21065 |  |
|  | Pangolin | Gx-P5L | Custom made by Genscript |
|  |  | GD-1 | Produced in-house in HEK293T cells as described previously (2) |
| 1A | Bat | Rs2018B | Produced in-house in HEK293T cells as described previously (2) |
|  |  | RsSHC014 |  |
|  |  | WIV-1 |  |
|  | SARS-CoV-1 | SARS-CoV-1 | Custom made by Genscript |

**Table S2.**
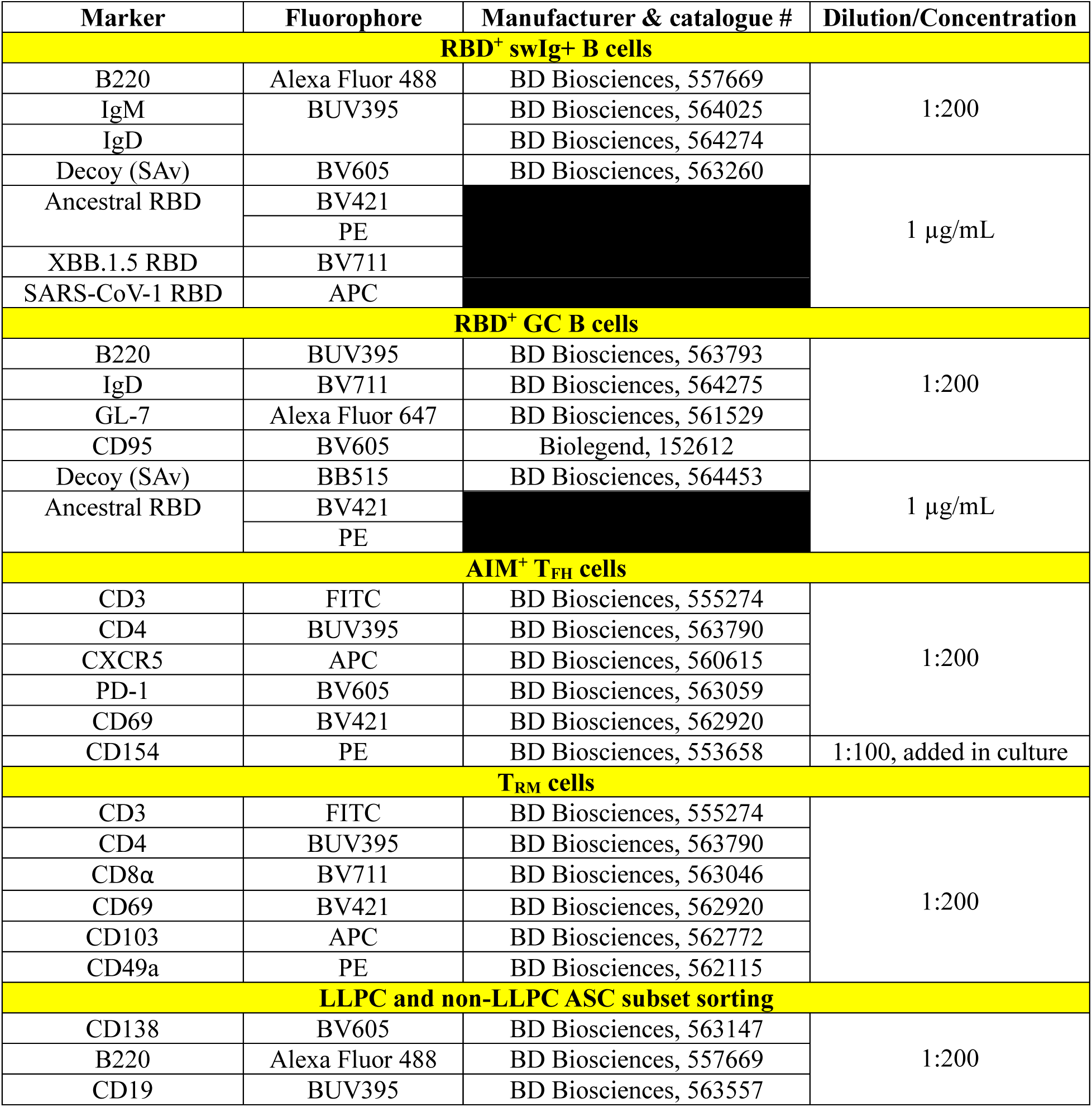
Antibodies used for flow cytometry staining of antigen-specific B cell subsets, AIM^+^ T_FH_ cells, T_RM_ cells, and sorting of LLPC and non-LLPC ASC subsets.

## REFERENCES

1. C. W. Tan, W. N. Chia, F. Zhu, B. E. Young, N. Chantasrisawad, S.-H. Hwa, A. Y.-Y. Yeoh, B. L. Lim, W. C. Yap, S. K. M. S. Pada, S. Y. Tan, W. Jantarabenjakul, L. K. Toh, S. Chen, J. Zhang, Y. Y. Mah, V. C.-W. Chen, M. I.-C. Chen, S. Wacharapluesadee, A. Sigal, O. Putcharoen, D. C. Lye, L.-F. Wang, SARS-CoV-2 Omicron variant emerged under immune selection. Nat Microbiol 7, 1756– 1761 (2022).

2. H. Guo, A. Li, T.-Y. Dong, J. Su, Y.-L. Yao, Y. Zhu, Z.-L. Shi, M. Letko, ACE2-Independent Bat Sarbecovirus Entry and Replication in Human and Bat Cells. mBio 13 (2022).

3. S. Maher, N. M. El Assaly, D. M. Aly, S. Atta, A. M. Fteah, H. Badawi, M. Y. Zahran, M. Kamel, Comparative study of neutralizing antibodies titers in response to different types of COVID-19 vaccines among a group of egyptian healthcare workers. Virol J 21, 277 (2024).

4. D. S. Khoury, D. Cromer, A. Reynaldi, T. E. Schlub, A. K. Wheatley, J. A. Juno, K. Subbarao, S. J. Kent, J. A. Triccas, M. P. Davenport, Neutralizing antibody levels are highly predictive of immune protection from symptomatic SARS-CoV-2 infection. Nat Med 27, 1205–1211 (2021).

5. R. R. Goel, M. M. Painter, S. A. Apostolidis, D. Mathew, W. Meng, A. M. Rosenfeld, K. A. Lundgreen, A. Reynaldi, D. S. Khoury, A. Pattekar, S. Gouma, L. Kuri-Cervantes, P. Hicks, S. Dysinger, A. Hicks, H. Sharma, S. Herring, S. Korte, A. E. Baxter, D. A. Oldridge, J. R. Giles, M. E. Weirick, C. M. McAllister, M. Awofolaju, N. Tanenbaum, E. M. Drapeau, J. Dougherty, S. Long, K. D’Andrea, J. T. Hamilton, M. McLaughlin, J. C. Williams, S. Adamski, O. Kuthuru, I. Frank, M. R. Betts, L. A. Vella, A. Grifoni, D. Weiskopf, A. Sette, S. E. Hensley, M. P. Davenport, P. Bates, E. T. Luning Prak, A. R. Greenplate, E. J. Wherry, mRNA vaccines induce durable immune memory to SARS-CoV-2 and variants of concern. Science (1979) 374 (2021).

6. E. Mitsi, M. O. Diniz, J. Reiné, A. M. Collins, R. E. Robinson, A. Hyder-Wright, M. Farrar, K. Liatsikos, J. Hamilton, O. Onyema, B. C. Urban, C. Solórzano, S. Belij-Rammerstorfer, E. Sheehan, T. Lambe, S. J. Draper, D. Weiskopf, A. Sette, M. K. Maini, D. M. Ferreira, Respiratory mucosal immune memory to SARS-CoV-2 after infection and vaccination. Nat Commun 14, 6815 (2023).

7. J. Tang, C. Zeng, T. M. Cox, C. Li, Y. M. Son, I. S. Cheon, Y. Wu, S. Behl, J. J. Taylor, R. Chakaraborty, A. J. Johnson, D. N. Shiavo, J. P. Utz, J. S. Reisenauer, D. E. Midthun, J. J. Mullon, E. S. Edell, M. G. Alameh, L. Borish, W. G. Teague, M. H. Kaplan, D. Weissman, R. Kern, H. Hu, R. Vassallo, S.-L. Liu, J. Sun, Respiratory mucosal immunity against SARS-CoV-2 after mRNA vaccination. Sci Immunol 7 (2022).

8. S. Ndeupen, Z. Qin, S. Jacobsen, A. Bouteau, H. Estanbouli, B. Z. Igyártó, The mRNA-LNP platform’s lipid nanoparticle component used in preclinical vaccine studies is highly inflammatory. iScience 24, 103479 (2021).

9. A. P. S. Rathore, A. L. St. John, Promises and challenges of mucosal COVID-19 vaccines. Vaccine 41, 4042–4049 (2023).

10. Y. Hu, Q. Wu, F. Chang, J. Yang, X. Zhang, Q. Wang, J. Chen, S. Teng, Y. Liu, X. Zheng, Y. Wang, R. Lu, D. Pan, Z. Liu, F. Liu, T. Xie, C. Wu, Y. Tang, F. Tang, J. Qian, H. Chen, W. Liu, Y.-P. Li, X. Qu, Broad cross neutralizing antibodies against sarbecoviruses generated by SARS-CoV-2 infection and vaccination in humans. NPJ Vaccines 9, 195 (2024).

11. Y. Pastor, N. Ghazzaui, A. Hammoudi, M. Centlivre, S. Cardinaud, Y. Levy, Refining the DC-targeting vaccination for preventing emerging infectious diseases. Front Immunol 13 (2022).

12. D. Yin, Y. Zhong, S. Ling, S. Lu, X. Wang, Z. Jiang, J. Wang, Y. Dai, X. Tian, Q. Huang, X. Wang, J. Chen, Z. Li, Y. Li, Z. Xu, H. Jiang, Y. Wu, Y. Shi, Q. Wang, J. Xu, W. Hong, H. Xue, H. Yang, Y. Zhang, L. Da, Z. Han, S. Tao, R. Dong, T. Ying, J. Hong, Y. Cai, Dendritic-cell-targeting virus-like particles as potent mRNA vaccine carriers. Nat Biomed Eng, doi: 10.1038/s41551-024-01208-4 (2024).

13. D. Horvath, N. Temperton, M. Mayora-Neto, K. Da Costa, D. Cantoni, R. Horlacher, A. Günther, A. Brosig, J. Morath, B. Jakobs, M. Groettrup, H. Hoschuetzky, J. Rohayem, J. ter Meulen, Novel intranasal vaccine targeting SARS-CoV-2 receptor binding domain to mucosal microfold cells and adjuvanted with TLR3 agonist Riboxxim^TM^ elicits strong antibody and T-cell responses in mice. Sci Rep 13, 4648 (2023).

14. S. Coléon, A. Wiedemann, M. Surénaud, C. Lacabaratz, S. Hue, M. Prague, M. Cervantes-Gonzalez, Z. Wang, J. Ellis, A. Sansoni, C. Pierini, Q. Bardin, M. Fabregue, S. Sharkaoui, P. Hoest, L. Dupaty, F. Picard, M. El Hajj, M. Centlivre, J. Ghosn, R. Thiébaut, S. Cardinaud, B. Malissen, G. Zurawski, A. Zarubica, S. M. Zurawski, V. Godot, Y. Lévy, Design, immunogenicity, and efficacy of a pan-sarbecovirus dendritic-cell targeting vaccine. EBioMedicine 80, 104062 (2022).

15. N. Y. Z. Cheang, K. Sen Tan, P. S. Tan, K. Purushotorma, W. C. Yap, K. M. Tullett, B. Y. L. Chua, A. Y.-Y. Yeoh, C. Q. H. Tan, X. Qian, H. Chen, D. J. W. Tay, I. Caminschi, Y. J. Tan, P. A. Macary, C. W. Tan, M. H. Lahoud, S. Alonso, Single-shot dendritic cell targeting SARS-CoV-2 vaccine candidate induces broad, durable and protective systemic and mucosal immunity in mice. Molecular Therapy 32, 2299–2315 (2024).

16. R. Kavishna, T. Y. Kang, M. Vacca, B. Y. L. Chua, H.-Y. Park, P. S. Tan, V. T. Chow, M. H. Lahoud, S. Alonso, A single-shot vaccine approach for the universal influenza A vaccine candidate M2e. Proceedings of the National Academy of Sciences 119 (2022).

17. H.-Y. Park, P. S. Tan, R. Kavishna, A. Ker, J. Lu, C. E. Z. Chan, B. J. Hanson, P. A. MacAry, I. Caminschi, K. Shortman, S. Alonso, M. H. Lahoud, Enhancing vaccine antibody responses by targeting Clec9A on dendritic cells. NPJ Vaccines 2, 31 (2017).

18. M. H. Lahoud, F. Ahmet, S. Kitsoulis, S. S. Wan, D. Vremec, C.-N. Lee, B. Phipson, W. Shi, G. K. Smyth, A. M. Lew, Y. Kato, S. N. Mueller, G. M. Davey, W. R. Heath, K. Shortman, I. Caminschi, Targeting Antigen to Mouse Dendritic Cells via Clec9A Induces Potent CD4 T Cell Responses Biased toward a Follicular Helper Phenotype. The Journal of Immunology 187, 842–850 (2011).

19. Y. Kato, A. Zaid, G. M. Davey, S. N. Mueller, S. L. Nutt, D. Zotos, D. M. Tarlinton, K. Shortman, M. H. Lahoud, W. R. Heath, I. Caminschi, Targeting Antigen to Clec9A Primes Follicular Th Cell Memory Responses Capable of Robust Recall. The Journal of Immunology 195, 1006–1014 (2015).

20. J. Li, F. Ahmet, L. C. Sullivan, A. G. Brooks, S. J. Kent, R. De Rose, A. M. Salazar, C. Reis e Sousa, K. Shortman, M. H. Lahoud, W. R. Heath, I. Caminschi, Antibodies targeting Clec9A promote strong humoral immunity without adjuvant in mice and non-human primates. Eur J Immunol 45, 854–864 (2015).

21. C.-W. Tan, W.-N. Chia, B. E. Young, F. Zhu, B.-L. Lim, W.-R. Sia, T.-L. Thein, M. I.-C. Chen, Y.-S. Leo, D. C. Lye, L.-F. Wang, Pan-Sarbecovirus Neutralizing Antibodies in BNT162b2-Immunized SARS-CoV-1 Survivors. New England Journal of Medicine 385, 1401–1406 (2021).

22. C. M. Freeman, J. L. Curtis, Lung Dendritic Cells: Shaping Immune Responses throughout Chronic Obstructive Pulmonary Disease Progression. Am J Respir Cell Mol Biol 56, 152–159 (2017).

23. D. Sichien, B. N. Lambrecht, M. Guilliams, C. L. Scott, Development of conventional dendritic cells: from common bone marrow progenitors to multiple subsets in peripheral tissues. Mucosal Immunol 10, 831–844 (2017).

24. Q. Wang, I. A. Mellis, Y. Guo, C. Gherasim, R. Valdez, A. Gordon, D. D. Ho, L. Liu, Robust SARS-CoV-2-neutralizing antibodies sustained through 6 months post XBB.1.5 mRNA vaccine booster. Cell Rep Med 5, 101701 (2024).

25. J. Tubiana, Y. Xiang, L. Fan, H. J. Wolfson, K. Chen, D. Schneidman-Duhovny, Y. Shi, Reduced B cell antigenicity of Omicron lowers host serologic response. Cell Rep 41, 111512 (2022).

26. X. Wang, S. Jiang, W. Ma, X. Li, K. Wei, F. Xie, C. Zhao, X. Zhao, S. Wang, C. Li, R. Qiao, Y. Cui, Y. Chen, J. Li, G. Cai, C. Liu, J. Yu, J. Li, Z. Hu, W. Zhang, S. Jiang, M. Li, Y. Zhang, P. Wang, Enhanced neutralization of SARS-CoV-2 variant BA.2.86 and XBB sub-lineages by a tetravalent COVID-19 vaccine booster. Cell Host Microbe 32, 25–34.e5 (2024).

27. S. Longet, A. Hargreaves, S. Healy, R. Brown, H. R. Hornsby, N. Meardon, T. Tipton, E. Barnes, S. Dunachie, C. J. A. Duncan, P. Klenerman, A. Richter, L. Turtle, T. I. de Silva, M. W. Carroll, mRNA vaccination drives differential mucosal neutralizing antibody profiles in naïve and SARS-CoV-2 previously-infected individuals. Front Immunol 13 (2022).

28. D. T. Hunt, J. L. Yates, K. E. Kulas, K. Carson, T. Lamson, V. Demarest, A. Furuya, K. Howard, M. Marchewka, R. Stone, H. Tucker, C. Warszycki, J. Yee, H. S. Yang, S. Racine-Brzostek, Z. Zhao, M. Ejemel, Q. Li, Y. Wang, S. Fernando, F. La Carpia, E. A. Hod, K. A. McDonough, W. T. Lee, COVID-19 Serology in New York State Using a Multiplex Microsphere Immunoassay. [Preprint] (2021). 10.1101/2021.05.12.21257125.

29. R. Wang, Y. Han, R. Zhang, J. Zhu, X. Nan, Y. Liu, Z. Yang, B. Zhou, J. Yu, Z. Lin, J. Li, P. Chen, Y. Wang, Y. Li, D. Liu, X. Shi, X. Wang, Q. Zhang, Y. R. Yang, T. Li, L. Zhang, Dissecting the intricacies of human antibody responses to SARS-CoV-1 and SARS-CoV-2 infection. Immunity 56, 2635–2649.e6 (2023).

30. Y. Feng, M. Yuan, J. M. Powers, M. Hu, J. E. Munt, P. S. Arunachalam, S. R. Leist, L. Bellusci, J. Kim, K. R. Sprouse, L. E. Adams, S. Sundaramurthy, X. Zhu, L. M. Shirreff, M. L. Mallory, T. D. Scobey, A. Moreno, D. T. O’Hagan, H. Kleanthous, F. J. Villinger, D. Veesler, N. P. King, M. S. Suthar, S. Khurana, R. S. Baric, I. A. Wilson, B. Pulendran, Broadly neutralizing antibodies against sarbecoviruses generated by immunization of macaques with an AS03-adjuvanted COVID-19 vaccine. Sci Transl Med 15 (2023).

31. X. Chen, L. Li, R. Du, Z. Wang, Y. Li, Y. Sun, R. Qin, H. Feng, L. Hu, X. Chen, M. Lu, X. Huang, L. Jiang, T. Zuo, B cells imprinted by ancestral SARS-CoV-2 develop pan-sarbecovirus neutralization in immune recalls. [Preprint] (2024). 10.1101/2024.10.13.618110.

32. Y. Hu, Q. Wu, F. Chang, J. Yang, X. Zhang, Q. Wang, J. Chen, S. Teng, Y. Liu, X. Zheng, Y. Wang, R. Lu, D. Pan, Z. Liu, F. Liu, T. Xie, C. Wu, Y. Tang, F. Tang, J. Qian, H. Chen, W. Liu, Y.-P. Li, X. Qu, Broad cross neutralizing antibodies against sarbecoviruses generated by SARS-CoV-2 infection and vaccination in humans. NPJ Vaccines 9, 195 (2024).

33. Y.-F. Hu, T. T.-T. Yuen, H.-R. Gong, B. Hu, J.-C. Hu, X.-S. Lin, L. Rong, C. L. Zhou, L.-L. Chen, X. Wang, C. Lei, T. Yau, I. F.-N. Hung, K. K.-W. To, K.-Y. Yuen, B.-Z. Zhang, H. Chu, J.-D. Huang, Rational design of a booster vaccine against COVID-19 based on antigenic distance. Cell Host Microbe 31, 1301–1316.e8 (2023).

34. C. W. Tan, S. A. Valkenburg, L. L. M. Poon, L.-F. Wang, Broad-spectrum pan-genus and pan-family virus vaccines. Cell Host Microbe 31, 902–916 (2023).

35. L. Peng, Z. Fang, P. A. Renauer, A. McNamara, J. J. Park, Q. Lin, X. Zhou, M. B. Dong, B. Zhu, H. Zhao, C. B. Wilen, S. Chen, Multiplexed LNP-mRNA vaccination against pathogenic coronavirus species. Cell Rep 40, 111160 (2022).

36. A. T. Tan, J. M. E. Lim, N. Le Bert, K. Kunasegaran, A. Chia, M. D. C. Qui, N. Tan, W. N. Chia, R. de Alwis, D. Ying, J. X. Y. Sim, E. E. Ooi, L.-F. Wang, M. I.-C. Chen, B. E. Young, Li Yang Hsu, J. G. H. Low, D. C. Lye, A. Bertoletti, Rapid measurement of SARS-CoV-2 spike T cells in whole blood from vaccinated and naturally infected individuals. Journal of Clinical Investigation 131 (2021).

37. P. J. Halfmann, K. Loeffler, A. Duffy, M. Kuroda, J. E. Yang, E. R. Wright, Y. Kawaoka, R. S. Kane, Broad protection against clade 1 sarbecoviruses after a single immunization with cocktail spike-protein-nanoparticle vaccine. Nat Commun 15, 1284 (2024).

38. E. Wang, A. A. Cohen, L. F. Caldera, J. R. Keeffe, A. V. Rorick, Y. M. Adia, P. N. P. Gnanapragasam, P. J. Bjorkman, A. K. Chakraborty, Designed mosaic nanoparticles enhance cross-reactive immune responses in mice. Cell, doi: 10.1016/j.cell.2024.12.015 (2025).

39. J. van Bergen, M. G. M. Camps, I. N. Pardieck, D. Veerkamp, W. Y. Leung, A. A. Leijs, S. K. Myeni, M. Kikkert, R. Arens, G. C. Zondag, F. Ossendorp, Multiantigen pan-sarbecovirus DNA vaccines generate protective T cell immune responses. JCI Insight 8 (2023).

40. D. C. Nguyen, I. T. Hentenaar, A. Morrison-Porter, D. Solano, N. S. Haddad, C. Castrillon, M. C. Runnstrom, P. A. Lamothe, J. Andrews, D. Roberts, S. Lonial, I. Sanz, F. E.-H. Lee, SARS-CoV-2-specific plasma cells are not durably established in the bone marrow long-lived compartment after mRNA vaccination. Nat Med 31, 235–244 (2025).

41. S. Crotty, T Follicular Helper Cell Biology: A Decade of Discovery and Diseases. Immunity 50, 1132–1148 (2019).

42. A. C. Olatunde, J. S. Hale, T. J. Lamb, Cytokine-skewed Tfh cells: functional consequences for B cell help. Trends Immunol 42, 536–550 (2021).

43. R. He, X. Zheng, J. Zhang, B. Liu, Q. Wang, Q. Wu, Z. Liu, F. Chang, Y. Hu, T. Xie, Y. Liu, J. Chen, J. Yang, S. Teng, R. Lu, D. Pan, Y. Wang, L. Peng, W. Huang, V. Terzieva, W. Liu, Y. Wang, Y.-P. Li, X. Qu, SARS-CoV-2 spike-specific TFH cells exhibit unique responses in infected and vaccinated individuals. Signal Transduct Target Ther 8, 393 (2023).

44. D. Ashour, P. Arampatzi, V. Pavlovic, K. U. Förstner, T. Kaisho, A. Beilhack, F. Erhard, M. B. Lutz, IL-12 from endogenous cDC1, and not vaccine DC, is required for Th1 induction. JCI Insight 5 (2020).

45. N. D. Bhattacharyya, C. G. Feng, Regulation of T Helper Cell Fate by TCR Signal Strength. Front Immunol 11 (2020).

46. J. P. Snook, C. Kim, M. A. Williams, TCR signal strength controls the differentiation of CD4+ effector and memory T cells. Sci Immunol 3 (2018).

47. G. Mantus, L. E. Nyhoff, V.-V. Edara, V. I. Zarnitsyna, C. R. Ciric, M. W. Flowers, C. Norwood, M. Ellis, L. Hussaini, K. E. Manning, K. Stephens, E. J. Anderson, R. Ahmed, M. S. Suthar, J. Wrammert, Pre-existing SARS-CoV-2 immunity influences potency, breadth, and durability of the humoral response to SARS-CoV-2 vaccination. Cell Rep Med 3, 100603 (2022).

48. Y. Chen, P. Tong, N. Whiteman, A. Sanjari Moghaddam, M. Zarghami, A. Zuiani, S. Habibi, A. Gautam, Keerti, C. Bi, T. Xiao, Y. Cai, B. Chen, D. Neuberg, D. R. Wesemann, Immune recall improves antibody durability and breadth to SARS-CoV-2 variants. Sci Immunol 7 (2022).

49. A. Tarke, P. Ramezani-Rad, T. Alves Pereira Neto, Y. Lee, V. Silva-Moraes, B. Goodwin, N. Bloom, L. Siddiqui, L. Avalos, A. Frazier, Z. Zhang, R. da Silva Antunes, J. Dan, S. Crotty, A. Grifoni, A. Sette, SARS-CoV-2 breakthrough infections enhance T cell response magnitude, breadth, and epitope repertoire. Cell Rep Med 5, 101583 (2024).

50. S. M. Lightman, A. Utley, K. P. Lee, Survival of Long-Lived Plasma Cells (LLPC): Piecing Together the Puzzle. Front Immunol 10 (2019).

51. R. Fraser, A. Orta-Resendiz, A. Mazein, D. H. Dockrell, Upper respiratory tract mucosal immunity for SARS-CoV-2 vaccines. Trends Mol Med 29, 255–267 (2023).

52. U. Marking, O. Bladh, S. Havervall, J. Svensson, N. Greilert-Norin, K. Aguilera, M. Kihlgren, A.-C. Salomonsson, M. Månsson, R. Gallini, C. Kriegholm, P. Bacchus, S. Hober, M. Gordon, K. Blom, A. Smed-Sörensen, M. Åberg, J. Klingström, C. Thålin, 7-month duration of SARS-CoV-2 mucosal immunoglobulin-A responses and protection. Lancet Infect Dis 23, 150–152 (2023).

53. R. Nakahashi-Ouchida, K. Fujihashi, Y. Kurashima, Y. Yuki, H. Kiyono, Nasal vaccines: solutions for respiratory infectious diseases. Trends Mol Med 29, 124–140 (2023).

54. X. Zhou, Y. Wu, Z. Zhu, C. Lu, C. Zhang, L. Zeng, F. Xie, L. Zhang, F. Zhou, Mucosal immune response in biology, disease prevention and treatment. Signal Transduct Target Ther 10, 7 (2025).

55. K. Chen, G. Magri, E. K. Grasset, A. Cerutti, Rethinking mucosal antibody responses: IgM, IgG and IgD join IgA. Nat Rev Immunol 20, 427–441 (2020).

56. G. Laghlali, M. J. Wiest, D. Karadag, P. Warang, J. J. O’Konek, L. A. Chang, S.-C. Park, V. Yan, M. Farazuddin, K. W. Janczak, A. García-Sastre, J. R. Baker, P. T. Wong, M. Schotsaert, Enhanced mucosal SARS-CoV-2 immunity after heterologous intramuscular mRNA prime/intranasal protein boost vaccination with a combination adjuvant. Molecular Therapy 32, 4448–4466 (2024).

57. M. Z. M. Zheng, L. M. Wakim, Tissue resident memory T cells in the respiratory tract. Mucosal Immunol 15, 379–388 (2022).

58. S. Yenyuwadee, J. L. Sanchez-Trincado Lopez, R. Shah, P. C. Rosato, V. A. Boussiotis, The evolving role of tissue-resident memory T cells in infections and cancer. Sci Adv 8 (2022).

59. K. P. J. M. van Gisbergen, K. D. Zens, C. Münz, T-cell memory in tissues. Eur J Immunol 51, 1310– 1324 (2021).

60. E. C. Reilly, M. Sportiello, K. L. Emo, A. M. Amitrano, R. Jha, A. B. R. Kumar, N. G. Laniewski, H. Yang, M. Kim, D. J. Topham, CD49a Identifies Polyfunctional Memory CD8 T Cell Subsets that Persist in the Lungs After Influenza Infection. Front Immunol 12 (2021).

61. D. K. J. Pieren, S. G. Kuguel, J. Rosado, A. G. Robles, J. Rey-Cano, C. Mancebo, J. Esperalba, V. Falcó, M. J. Buzón, M. Genescà, Limited induction of polyfunctional lung-resident memory T cells against SARS-CoV-2 by mRNA vaccination compared to infection. Nat Commun 14, 1887 (2023).

62. P. B. Gilbert, D. C. Montefiori, A. B. McDermott, Y. Fong, D. Benkeser, W. Deng, H. Zhou, C. R. Houchens, K. Martins, L. Jayashankar, F. Castellino, B. Flach, B. C. Lin, S. O’Connell, C. McDanal, A. Eaton, M. Sarzotti-Kelsoe, Y. Lu, C. Yu, B. Borate, L. W. P. van der Laan, N. S. Hejazi, C. Huynh, J. Miller, H. M. El Sahly, L. R. Baden, M. Baron, L. De La Cruz, C. Gay, S. Kalams, C. F. Kelley, M. P. Andrasik, J. G. Kublin, L. Corey, K. M. Neuzil, L. N. Carpp, R. Pajon, D. Follmann, R. O. Donis, R. A. Koup, Immune correlates analysis of the mRNA-1273 COVID-19 vaccine efficacy clinical trial. Science (1979) 375, 43–50 (2022).

63. D. S. Khoury, D. Cromer, A. Reynaldi, T. E. Schlub, A. K. Wheatley, J. A. Juno, K. Subbarao, S. J. Kent, J. A. Triccas, M. P. Davenport, Neutralizing antibody levels are highly predictive of immune protection from symptomatic SARS-CoV-2 infection. Nat Med 27, 1205–1211 (2021).

64. A. Wajnberg, F. Amanat, A. Firpo, D. R. Altman, M. J. Bailey, M. Mansour, M. McMahon, P. Meade, D. R. Mendu, K. Muellers, D. Stadlbauer, K. Stone, S. Strohmeier, V. Simon, J. Aberg, D. L. Reich, F. Krammer, C. Cordon-Cardo, Robust neutralizing antibodies to SARS-CoV-2 infection persist for months. Science (1979) 370, 1227–1230 (2020).

65. J. M. Knisely, L. E. Buyon, R. Mandt, R. Farkas, S. Balasingam, K. Bok, U. J. Buchholz, M. P. D’Souza, J. L. Gordon, D. F. L. King, T. T. Le, W. W. Leitner, R. A. Seder, A. Togias, S. Tollefsen, D. W. Vaughn, D. N. Wolfe, K. L. Taylor, A. S. Fauci, Mucosal vaccines for SARS-CoV-2: scientific gaps and opportunities—workshop report. NPJ Vaccines 8, 53 (2023).

66. J. Jonny, T. A. Putranto, M. L. Yana, E. C. Sitepu, R. Irfon, B. P. Ramadhani, M. A. U. Sofro, Y. M. Nency, E. S. Lestari, R. Triwardhani, Mujahidah, R. K. Sari, N. A. Soetojo, Safety and efficacy of dendritic cell vaccine for COVID-19 prevention after 1-Year follow-up: phase I and II clinical trial final result. Front Immunol 14 (2023).

67. G. I. Nistor, R. O. Dillman, R. M. Robles, J. L. Langford, A. J. Poole, M. A. U. Sofro, Y. M. Nency, J. Jonny, M. L. Yana, M. Karyana, E. S. Lestari, R. Triwardhani, M. Mujahidah, R. K. Sari, N. A. Soetojo, D. Wibisono, D. Tjen, T. Ikrar, G. Sarkissian, H. Winarta, T. A. Putranto, H. S. Keirstead, A personal COVID-19 dendritic cell vaccine made at point-of-care: Feasibility, safety, and antigen-specific cellular immune responses. Hum Vaccin Immunother 18 (2022).

68. T. Granot, T. Senda, D. J. Carpenter, N. Matsuoka, J. Weiner, C. L. Gordon, M. Miron, B. V. Kumar, A. Griesemer, S.-H. Ho, H. Lerner, J. J. C. Thome, T. Connors, B. Reizis, D. L. Farber, Dendritic Cells Display Subset and Tissue-Specific Maturation Dynamics over Human Life. Immunity 46, 504–515 (2017).

69. K. M. Tullett, I. M. Leal Rojas, Y. Minoda, P. S. Tan, J.-G. Zhang, C. Smith, R. Khanna, K. Shortman, I. Caminschi, M. H. Lahoud, K. J. Radford, Targeting CLEC9A delivers antigen to human CD141+ DC for CD4+ and CD8+T cell recognition. JCI Insight 1 (2016).

